# Strain-aware assembly of genomes from mixed samples using flow variation graphs

**DOI:** 10.1101/645721

**Authors:** Jasmijn A. Baaijens, Leen Stougie, Alexander Schönhuth

## Abstract

The goal of strain-aware genome assembly is to reconstruct all individual haplotypes from a mixed sample at the strain level and to provide abundance estimates for the strains. Given that the use of a reference genome can introduce significant biases, de novo approaches are most suitable for this task. So far, reference-genome-independent assemblers have been shown to reconstruct haplotypes for mixed samples of limited complexity and genomes not exceeding 10000 bp in length.

Here, we present VG-Flow, a de novo approach that enables full-length haplotype reconstruction from pre-assembled contigs of complex mixed samples. Our method increases contiguity of the input assembly and, at the same time, it performs haplotype abundance estimation. VG-Flow is the first approach to require polynomial, and not exponential runtime in terms of the underlying graphs. Since runtime increases only linearly in the length of the genomes in practice, it enables the reconstruction also of genomes that are longer by orders of magnitude, thereby establishing the first de novo solution to strain-aware full-length genome assembly applicable to bacterial sized genomes.

VG-Flow is based on the *flow variation graph* as a novel concept that both captures all diversity present in the sample and enables to cast the central contig abundance estimation problem as a flow-like, polynomial time solvable optimization problem. As a consequence, we are in position to compute maximal-length haplotypes in terms of decomposing the resulting flow efficiently using a greedy algorithm, and obtain accurate frequency estimates for the reconstructed haplotypes through linear programming techniques.

Benchmarking experiments show that our method outperforms state-of-the-art approaches on mixed samples from short genomes in terms of assembly accuracy as well as abundance estimation. Experiments on longer, bacterial sized genomes demonstrate that VG-Flow is the only current approach that can reconstruct full-length haplotypes from mixed samples at the strain level in human-affordable runtime.

## Introduction

Sequencing datasets relating to bacterial or viral samples, such as metagenomes or viral quasispecies [1] as most prominent examples, contain mixtures of genomes that do not only differ at the level of species, but at the level of the individual bacterial or viral strains. Since relevant properties such as escape from medical treatment or host immune response happen at the strain, and not the species level [2, 3], strain-aware reconstruction of genomes from mixed samples is an important, urgent issue. The challenge in this is to distinguish between strain-level genetic variation, which is comparatively minor, and to estimate the relative abundances of the individual strains in the mix, at sufficient accuracy. Thereby, a particularly important point is that highly abundant, hence dominant strains may mask strains of lower abundance, where low-abundance strains are often most relevant factors when considering resistance to treatment, or other clinical concerns.

Because we emphasize the genetic variation of the individual (strain-level) genomes, we refer to the individual genomes as haplotypes in the following, and realize that the challenge more generally can be referred to as *strain-level haplotype-aware genome assembly from mixed samples*. Related approaches presented so far do either depend on reference genomes [4, 5], hence tend to suffer from considerable biases during strain-level reconstruction, while reference-independent, de novo approaches are applicable to mixed samples of genomes not exceeding 10000 bp in length [6, 7, 8]. This renders such approaches applicable to viral quasispecies in the first place, but one cannot use them for bacterial sized genomes. In experiments presented here, we notice that even reference-dependent approaches tend to struggle with bacterial sized genomes.

In this paper, we present VG-Flow as a method that overcomes the above-mentioned challenges. VG-Flow operates in de novo mode, hence does not suffer from any reference-induced biases. The algorithm performs full-length haplotype reconstruction from pre-assembled contigs, thereby increasing assembly contiguity while maintaining low error rates. Crucially, VG-Flow overcomes a computational bottleneck in our previous approach that yielded computational runtimes exponential in the number of vertices and edges of the underlying graphs, and, as a consequence, exponential in the length of the underlying genomes [7]. We propose a solution that replaces the exponential step in [7] by a polynomial algorithm. Hence, the overall runtime of our approach is polynomial in the genome size. This enables processing of genomes longer than viral genomes on orders of magnitude, such as bacterial genomes.

The methodical novelty that underlies VG-Flow’s advances is to derive *flow variation graphs* from the (common) variation graphs that one constructs from the input contigs. General variation graphs [9, 10] derived from input contigs had been presented in earlier work as a general means for overcoming linear reference induced biases and aiming at the reconstruction of full-length strain-level haplotypes [7]. The advances of VG-Flow are due to the construction of the corresponding flow variation graphs, which are a novel concept in general, and to cast the relevant computational problems in terms of these flow variation graphs in particular, which renders the problems polynomial-time solvable for the first time.

In this, first and foremost, VG-Flow presents the first comprehensive solution to assembling haplotypes from mixed samples at the strain level, also for longer genomes and samples of considerably increased complexity. It is further relevant to observe that VG-Flow also leads to improvements when processing shorter genomes, such as viral quasispecies; because of particularly high mutation rates, these data sets had still been leaving researchers with unresolved issues. Thereby, VG-Flow does not only lead to improvements in terms of runtime, but also in terms of assembly quality.

Note, finally, that reconstructing all of the individual haplotypes present in a metagenome, that is *de novo, strain-aware metagenome assembly*, is challenging also because of various other reasons, all of which require specialized tools and further methodological progress [11]. In that context, strain-aware metagenome assembly still casts considerable challenges, the majority of which are due to repeats and coverage fluctuations. Here, this means that the construction of common variation graphs from metagenomes may still be cumbersome. When taking all steps into an overall account, VG-Flow does not yet solve the issue of de novo strain-aware full-length metagenome assembly, but offers an important and so far missing, fundamental building block.

Benchmarking experiments finally demonstrate the superiority of VG-Flow with respect to the above-mentioned, so far unresolved issues in comparison with state-of-the-art approaches, considering both tools specializing in (both reference based and de novo) strain-aware assembly [4, 6, 7, 12, 13, 14, 15, 8, 16] and in (haplotype-aware) de novo genome assembly [17, 18, 19, 20, 21], which encompasses all applicable approaches.

Please also see ‘Further Related Work’ in the Supplement for a list of problems and solutions that further relate to VG-Flow, such as theory referring to flow decomposition and work on RNA transcript assembly [22, 23, 24, 25, 26].

## Methods

We present VG-Flow, a new approach to haplotype-aware genome assembly from mixed samples. This algorithm takes as input a data set of next-generation sequencing reads and a collection of strain-specific contigs assembled from the data (which can be obtained using already available tools specializing in the generation of haplotype-aware contigs; see also ‘Remark on generation of haplotype-aware contigs’ in the Supplementary Materials), and produces full-length haplotypes and corresponding abundance estimates. Our method is centered on estimating contig abundances in a *contig variation graph*, a graph that captures all quasispecies diversity present in the contigs [9, 10]. We build a *flow network* (also referred to as *flow variation graph*) to accompany the variation graph and estimate contig abundances by solving a flow-like optimization problem: variables represent flow values on the edges of the flow variation graph and we impose flow constraints, while the objective function evaluates the difference between estimated contig abundances and read coverage for every node in the variation graph. This objective function is convex, which renders the flow problem polynomial time solvable [27]. Note that de novo assembly of strain-specific contigs can be performed using various tools (e.g. [6, 8, 17, 18], depending on the application). The output of VG-Flow consists of maximal length haplotypes along with relative abundance estimates for each of these sequences. In addition, the algorithm yields abundance estimates for each of the input contigs. In earlier work, the steps after the construction of the contig variation graph overall required runtime exponential in the number of vertices and edges of that graph. Since numbers of vertices and edges scale linearly with genome size in practice, this prohibited to process mixtures of genomes longer than 10 kbp. The theoretical improvement here is that all steps following the construction of the contig variation graph now only require runtime polynomial in the vertices and edges. In our experiments, we further demonstrate that VG-flow scales approximately linearly in genome size in practice (Supplemental Figures 1 and 2), which allows to process mixtures of genomes that are longer on orders of magnitude. See paragraph ‘Theoretical Runtime Analysis’ for a more detailed discussion.

### Algorithm overview

Our approach consists of five steps, as depicted in Figure 1:

1. We construct a *contig variation graph V G*_*C*_ by performing Multiple Sequence Alignment (MSA) on the input sequences. Node abundances are obtained by mapping sequencing reads to the variation graph.
2. We build a *flow variation graph FV G* using *V G*_*C*_.
3. We define and solve a flow-like optimization problem on *FV G* to obtain contig abundance estimates.
4. We generate a set of candidate haplotypes *P*_cand_ based on the estimated contig abundances through multiple greedy heuristics.
5. We obtain a selection of haplotypes *H* from *P*_cand_ by solving another linear optimization problem, defined on *V G*_*C*_. The solution to this problem presents estimates for the relative abundances of all candidate haplotypes in *P*_cand_, thereby eliminating any false haplotypes.

**Figure 1:**
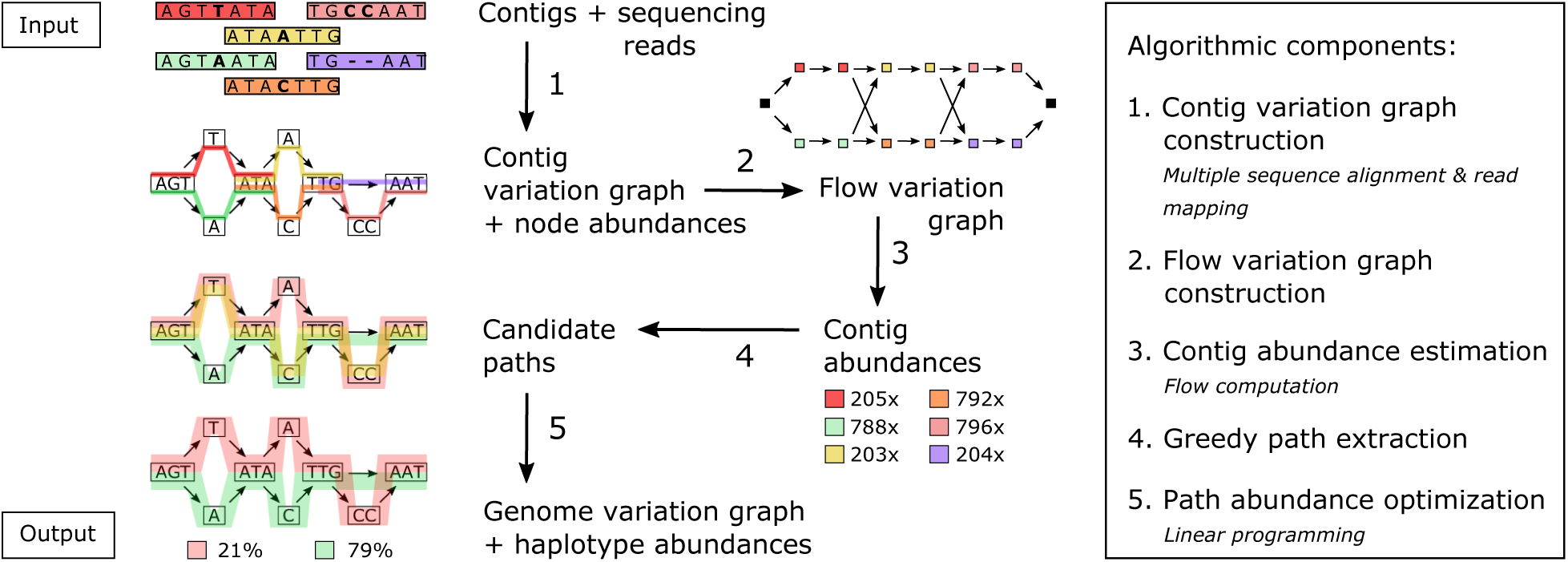
Algorithm overview.

The final output is presented as a *genome variation graph V G*_*H*_ capturing the haplotypes in *H*, along with the estimated relative abundances. While the already efficient or practially feasible steps (1) and (5) correspond to prior work [7], steps (2), (3) and (4) are novel, and replace the exponential-runtime procedure presented earlier [7]. As benefits, VG-flow can process mixtures that are both more complex and refer to genomes longer on orders of magnitude, now clearly including bacterial sized genomes.

### Variation graphs

Variation graphs are mathematical structures that capture genetic variation between haplotypes in a population [9, 10]. These graphs provide a compact representation of a collection of input sequences by collapsing all shared subsequences between haplotypes.

#### Definition

Let *S* be a collection of sequences. We define the variation graph *V G*(*S*) as a tuple (*V, E, P, a*). The nodes *v* ∈ *V* store sequences seq(*v*) of nucleotides (of arbitrary length) which appear as a substring of some *s* ∈ *S*. The edges (*v*_1_, *v*_2_) ∈ *E* indicate that the concatenation seq(*v*_1_)seq(*v*_2_) also appears as a substring of some *s* ∈ *S*. In addition to nodes *V* and edges *E*, a variation graph stores a set of paths *P* representing the input sequences: for every *s* ∈ *S* there is a path *p* ∈ *P* (i.e. a list of nodes, linked by edges) such that the concatenation of node sequences equals *s*. Finally, we store path abundances using an abundance function *a* : *P* → ℝ which assigns an absolute abundance value to each path in *P*.

#### Approach

Following [7], we distinguish between two types of variation graphs: *contig variation graphs* and *genome variation graphs*. Let *C* be a set of pre-assembled contigs and let *H* be the collection of haplotypes we aim to reconstruct. The contig variation graph *V G*(*C*) = (*V*_*C*_, *E*_*C*_, *P*_*C*_, *a*_*C*_) organizes the genetic variation that is present in the input contigs and the abundance function *a*_*C*_ gives contig abundance values for every input contig. The genome variation graph *V G*(*H*) = (*V*_*H*_, *E*_*H*_, *P*_*H*_, *a*_*H*_) stores the haplotypes within a population and the abundance function computes haplotype abundances. Constructing a genome variation graph is the goal of our method; the key idea is to use the contig variation graph to get there.

### Contig variation graph construction

We construct a contig variation graph from pre-assembled contigs *C* adopting techniques from prior work [7]. This entails **(1)** multiple sequence alignment, for which we run vg msga [10] on the input contigs, representing the alignment as a graph (*V, E, P*), **(2)** compactifying (*V, E, P*) into (*V*_*C*_, *E*_*C*_, *P*_*C*_) by contracting non-branching paths into single nodes, and **(3)** computing node abundances, reflecting the read coverage of the sequence patches (as average of the coverage of the individual bases) corresponding to the nodes using vg map [10].

We aim at computing full-length path coverages, as a function *a*_*C*_ : *P*_*C*_ → ℝ, from the node abundances. We do this in two steps: first, we compute *contig abundances* by means of an adaptation of a minimum-cost flow problem, from which we finally are in position to compute *a*_*C*_.

### Flow variation graph construction

We build a flow variation graph *FV G* = (*V, E, c, d*), which is a flow network that allows us to compute contig abundances by solving a linear optimization problem that is a variant of the minimum-cost flow problem. Network flows are defined on directed graphs, where every edge has a given capacity and receives a certain amount of flow [28]. A flow variation graph has a *source* node and a *sink* node, which have only incoming and outgoing flow, respectively. For all other nodes, the amount of incoming flow must always equal the amount of outgoing flow, so-called flow conservation. We define our graph as described below and illustrated in Figure 2.

**Figure 2:**
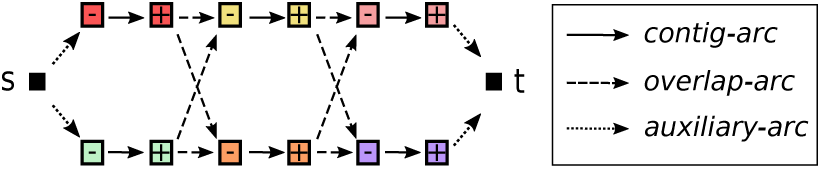
A flow variation graph with source (*s*), sink (*t*), vertices 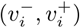, contig-arcs, overlap-arcs, and auxiliary-arcs.

**Nodes (***V* ). We start by creating a source *s* and a sink *t*. Then, we introduce two vertices for every contig *c*_*i*_ ∈ *C*, thus obtaining the vertex set 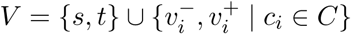.

**Edges (***E*). We introduce directed edges (arcs) of three types: *contig-arcs, overlap-arcs*, and *auxiliary-arcs*. For each contig *c*_*i*_ we add a contig-arc 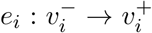. For each pair of contigs *c*_*i*_, *c*_*j*_ let *p*_*i*_, *p*_*j*_ ∈ *P*_*C*_ be the corresponding paths in *V G*(*C*). We add an overlap-arc *e*_*ij*_ to *FV G* from vertex 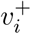 to vertex 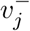 if a suffix of the path *p*_*i*_ has a non-conflicting, non-empty overlap with a prefix of the path *p*_*j*_. In other words, the sequences of *c*_*i*_ and *c*_*j*_ are identical on their overlap in the contig variation graph. Finally, we add auxiliary-arcs 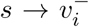 for any 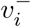 which has no incoming overlap-arcs, and auxiliary-arcs 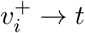 for any 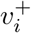 which has no outgoing overlap-arcs.

**Capacities (***c*). All edges have infinite capacity.

**Costs (***d*). To every edge *e* ∈ *E*, we assign a cost *d*_*e*_ where 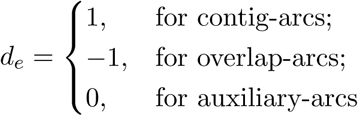.

The intuition is that *s* − *t* paths in *FV G* reflect haplotypes, while flow along the edges reflects accumulated (because several haplotypes may pass through identical edges) haplotype abundances. By defining and solving a min-cost flow problem, we can determine contig abundances that are optimal in terms of being compatible with the node abundances.

### Contig abundance computation

The problem of estimating contig abundances has applications in metagenomics [29] and RNA transcript assembly [30]. Existing methods make use of read mapping, either to a reference genome [31] or to the contigs [32, 33]. Although looking straightforward at first glance, accurately estimating contig abundances can be involved because of overlapping contigs and ambiguous read alignments. We solve this problem here by casting it as a flow-like optimization problem.

#### Problem formulation

Candidate haplotypes in the contig variation graph *V G*_*C*_ can be obtained by concatenating overlapping contig subpaths. Therefore, any maximal length path in the contig variation graph corresponds to an *s*-*t* path in *FV G*. We denote by *δ*^+^(*v*) and *δ*^−^(*v*) the set of arcs, respectively, entering and leaving *v* ∈ *V*. Recall that *V*_*C*_ denotes the set of nodes in *V G*_*C*_ and let 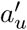 denote the abundance of node *u* ∈ *V*_*C*_, see above. For a node *u* ∈ *V*_*C*_ and edge *e* ∈ *E*, we write *u* ∈ *e* if the contig (or overlap) associated with the contig-arc (or overlap-arc) passes through node *u* in the contig variation graph. We define the following flow problem, in which the variables *x*_*e*_ decide the amount of flow going through arc *e* ∈ *E*:

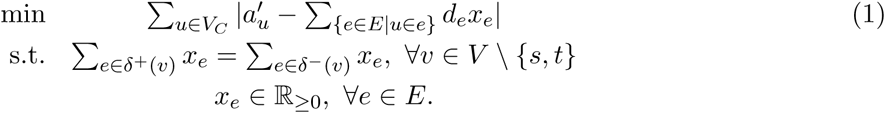

#### Motivation

The objective function reflects the sum of the absolute differences of the node abundances 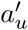 and the sum of the abundances of the contigs passing through the respective nodes ∑_{*e*∈*E*|*u*∈*e*}_ *d*_*e*_*x*_*e*_. Thereby, the edge costs *d*_*e*_ = −1 referring to overlap arcs prevent to double-count contig abundances for overlapping contigs when determining the flow through the flow variation graph. Given a solution to this optimization problem, the flow values on the contig-arcs reflect contig abundance estimates.

#### Solution

The objective function is convex in the flow-variables *x*_*e*_ (Supplementary Material, Lemma 3.1), so we have the problem of minimizing a convex function over a set of linear constraints, which is polynomial time solvable [27]. An evaluation of the accuracy of the contig abundance estimates, reflected by the contig-arc flow values, is presented in the Supplementary Materials.

### Greedy path extraction

The final steps are concerned with decomposing the flow (which provides flow values for all, and not just contig arcs) into *s*-*t* paths, which are supposed to reflect full-length haplotypes, and assign meaningful path abundances to the components. In this, we are interested in a parsimonious solution (as a general principle that agrees with biological reality), reflected by a decomposition of the flow into a minimal number of *s*-*t* paths. This amounts to seeking a solution of the minimum path flow decomposition problem, which unlike the flow decomposition problem without the minimality constraint [28], is NP-hard [34, 35]. Since various approximation algorithms referring to this problem (e.g. [36, 37]) were not able to handle even our smallest datasets (2 haplotypes of length 2.5 kbp), we resorted to greedy heuristics based, efficient means for obtaining a set of haplotypes from the given flow solution, thereby guaranteeing runtime to be polynomial in the size of the graph [35]. In this, we consider three optimality criteria for greedily picking *s* − *t* paths: maximum capacity, minimum capacity, and shortest paths. We update the flow by subtracting the greedily selected optimal path from the overall network flow; note that only flow on contig-arcs is considered, because modifications on overlap- or auxiliary arcs would impose restrictions on the usage of contigs. See Algorithm 1 for pseudo-code.

#### Algorithm 1 Greedy path extraction given a flow solution

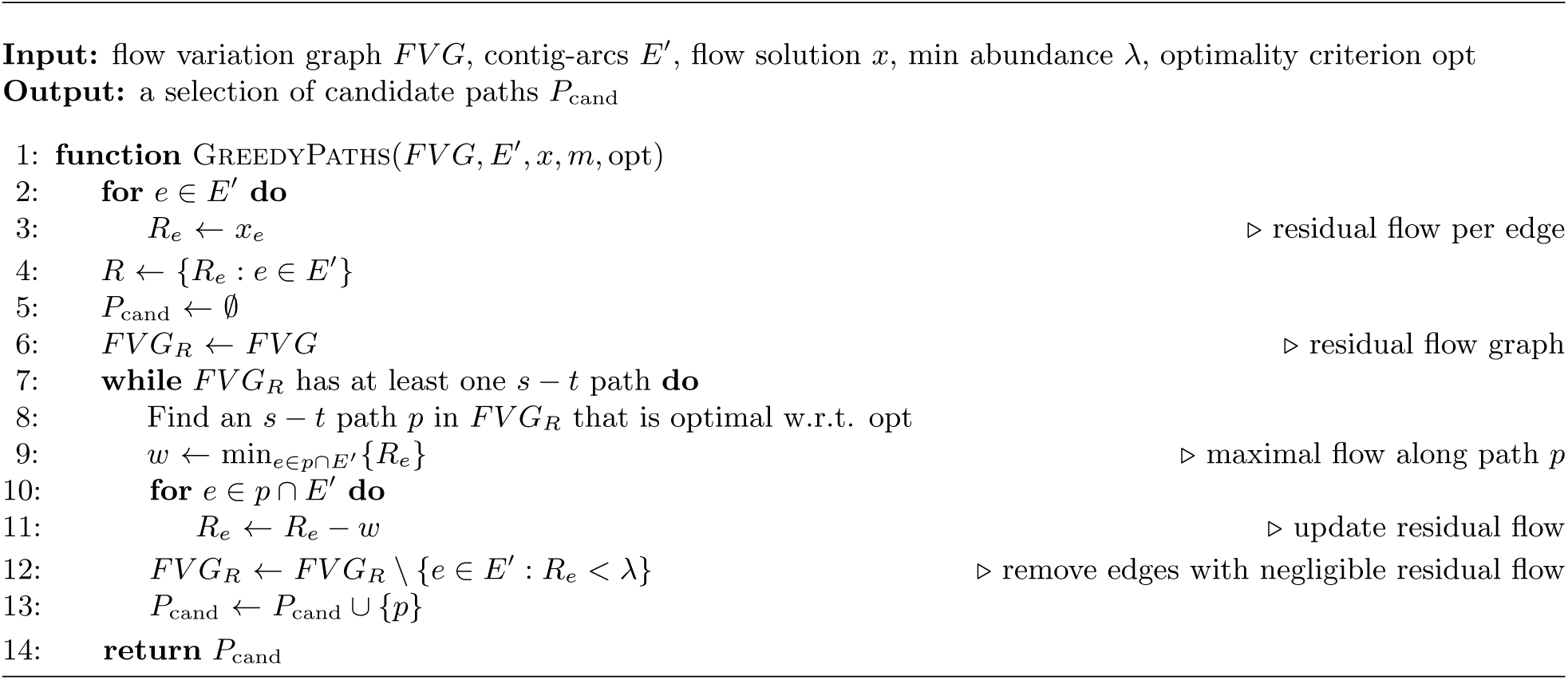

While the maximum capacity optimality criterion generally leads to very reliable paths, it tends to neglect true low-frequency haplotypes, which in turn are easily selected by the minimum capacity criterion. Because we do not know the composition of the mixture in general, we combine the results of all three heuristics employed. This strategy follows work that proves that this yields efficient, high quality approximations [38, 39]. See the Supplementary Material for an evaluation of all strategies tested.

### Path abundance optimization

Given a collection of candidate haplotypes *P*_cand_ in the form of paths through the contig variation graph, the only task remaining is to compute relative abundances for these haplotypes. Although the greedy path extraction algorithm produces preliminary path abundance estimates, these can be improved by the following linear programming approach, patterned according to [7], while here not dealing with an exponential number of candidate paths.

#### Problem formulation

Let 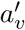 denote the abundance of node *v* ∈ *V*_*C*_, which was computed from the read alignments to *V G*_*C*_. We define variables *x*_*p*_ ∈ ℝ_≥0_ for *p* ∈ *P*_cand_, representing the estimated abundance for haplotype *p*, and consider the following optimization problem:

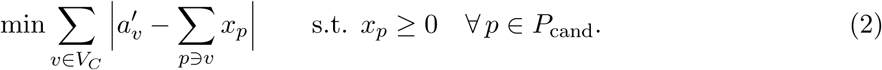

The objective function is similar in spirit to the one from (1), but only now we consider the absolute difference between the node abundance value and abundance estimates for full-length *s*-*t* paths, that is, all haplotypes passing through this node. This is a convex programming formulation, which can be linearized and solved using an LP solver, hence polynomial-time solvable in theory [27].

### Genome variation graph construction

Given the candidate haplotypes *P*_cand_ and the abundance estimates *x*_*p*_ for *p* ∈ *P*_cand_, we obtain a final selection of haplotypes *H* = {*p* ∈ *P*_cand_ : *x*_*p*_ ≥ *λ*}. Here, *λ* is a user-defined minimal path abundance, by default set to 1% of total sequencing depth. Given *H*, we can transform the contig variation graph *V G*_*C*_ into the genome variation graph *V G*_*H*_, a complete representation of the viral quasispecies.

### Theoretical runtime analysis

The first step of VG-Flow is to construct the contig variation graph. This requires multiple sequence alignment, which is quadratic in the number of pre-assembled contigs [40]. Since the vertices and edges of the flow variation graph scale linearly in its counterparts of the contig variation graph, construction of the flow variation graph is a polynomial-time procedure. Solving the contig abundance problem refers to computing the flow in the flow variation graph, which refers to minimizing a convex function over a set of linear constraints, hence can be done in time polynomial in the vertices and edges of the flow variation graph [27]. The greedy path extraction step refers to decomposing a flow into a set of s-t-paths while provably guaranteeing not to exceed upper bounds, linear in the edges, on the number of such paths [35]. Therefore, the runtime of the greedy algorithm is linear in the size of the graph as well. The last step, *path abundance optimization*, refers to a convex program referring to vertices and edges of the contig variation graph *V G*_*C*_, hence requires time polynomial in the vertices and edges of *V G*_*C*_ [27]. Constructing the genome variation graph, finally, is a simple filtering procedure that can be performed in time linear in the number of paths, which, by the properties of the greedy algorithm, were polynomial in terms of vertices and edges of the flow variation graph. This establishes polynomial runtime overall.

## Results

We perform benchmarking experiments where we compare VG-Flow to existing methods for fulllength viral quasispecies reconstruction. In these experiments, we make use of the specialized de novo viral quasipecies assembler SAVAGE [6] for generating a set of strain-specific contigs. We compare performance of VG-Flow to Virus-VG [7], another de novo approach, and to reference-guided viral quasispecies reconstruction tools PredictHaplo [12] and ShoRAH [4]. More recent viral quasispecies assemblers aBayesQR [14], QSdpR [15], and PEHaplo [8] focus on reconstruction of relatively short genomic regions; [14] and [15] are reference-guided and [8] is de novo, but none of these could finish processing our full-length quasispecies data sets at ultra-deep coverage within 500 hours. We provide the reference-guided methods with a *consensus reference genome* obtained by running VICUNA [41] on the data set, which generates a single sequence to represent the data. This procedure simulates a de novo setting where the viral agent and its genome may be unknown. Moreover, the consensus reference sequence may be a more accurate representation of the data set under consideration than the standard reference genomes available.

### Data simulation

All synthetic data sets were generated using the software SimSeq[42] to simulate Illumina MiSeq reads (2×250 bp) from the genome of interest. In order to obtain realistic sequencing error profiles, we used the MiSeq error profile provided with the software. The genomes used for each data set are listed in the Supplementary Material.

### Assembly evaluation

We evaluate all assemblies by comparing the assembled contigs to the ground truth sequences using QUAST [43]. For each assembly, we report the number of contigs, percent target genomes covered, NG50, and error rate. Target genome coverage is defined as the percentage of aligned bases in the true haplotypes, where a base is considered aligned if there is at least one contig with at least one alignment to this base. The NG50 reflects assembly contiguity and is defined as the length for which all contigs in the assembly of at least this length together add up to at least half of the total target length. Error rates reflect the number of errors relative to the genome size, calculated as the sum of mismatch rate, indel rate, and N-rate (ambiguous bases).

### Availability of data and material

Software and analysis scripts are publicly available at https://bitbucket.org/jbaaijens/vg-flow. The synthetic benchmarking datasets analysed during the current study are available at https://bitbucket.org/jbaaijens/savage-benchmarks.

## VG-Flow outperforms existing tools

### Short Genomes (Viral Quasispecies)

We evaluate performance of VG-Flow on three challenging synthetic viral quasispecies data sets from [7], capturing HCV, ZIKV, and Poliovirus mixtures. The de novo approaches VG-Flow and Virus-VG both use the contigs obtained with SAVAGE as input. We observe that both methods produce full-length haplotypes for all simulated data sets, with much higher assembly NG50 values than SAVAGE. The improved assembly contiguity comes with a decrease in target coverage compared to SAVAGE contigs for some of the data sets. This may be explained by the fact that contigs with abundance estimates below a user-defined threshold *λ* are removed from *P*_cand_ to ensure correctness. As a result, VG-Flow contigs have only slightly higher error rates than the SAVAGE contigs (Table 1a). Moreover, VG-Flow builds contigs with even lower error rates than Virus-VG (0.108% versus 0.115% on ZIKV data and 0.036% versus 0.064% on Poliovirus data for VG-Flow and Virus-VG, respectively). On the Poliovirus data set we do not only observe a lower error rate for VG-Flow compared to Virus-VG, but also a higher target coverage (90.2% for VG-Flow versus 80.7% for Virus-VG).

**Table 1:** Assembly results on simulated Illumina MiSeq data. ER = Error Rate (N’s + mismatches + indels). NG50 = the length for which all contigs in the assembly of at least this length together add up to at least half of the total target length. The high, overestimated number of full-length strains for *L*=100kbp,200kbp is due to ‘ambiguity islands’ among the different strains, an issue that can only be overcome by making use of longer reads. *If contigs are full-length, this number reflects the estimated number of strains in the quasispecies.

|  | # contigs* | target (%) | NG50 | ER(%) |  | # contigs* | target (%) | NG50 | ER(%) |
| --- | --- | --- | --- | --- | --- | --- | --- | --- | --- |
| *10-strain HCV* |  |  |  |  | *L = 10 kbp* |  |  |  |  |
| SAVAGE | 26 | 99.4 | 8964 | 0.001 | SAVAGE | 428 | 95.7 | 737 | 0.003 |
| Virus-VG | 10 | 99.3 | 9203 | 0.001 | Virus-VG | - | - | - | - |
| **VG-Flow** | 10 | 99.3 | 9203 | 0.001 | **VG-Flow** | 43 | 80.6 | 8908 | 0.485 |
| PredictHaplo | 9 | 73.8 | 7608 | 0.059 | PredictHaplo | 2 | 24.6 | - | 1.009 |
| ShoRAH | 639 | 56.9 | 7570 | 4.294 | ShoRAH | 84 | 85.5 | 9866 | 1.359 |
| *15-strain ZIKV* |  |  |  |  | *L = 100 kbp* |  |  |  |  |
| SAVAGE | 100 | 98.8 | 3801 | 0.023 | SAVAGE | 1059 | 80.6 | 1411 | 0.017 |
| Virus-VG | 20 | 92.8 | 10210 | 0.115 | Virus-VG | - | - | - | - |
| **VG-Flow** | 21 | 92.8 | 10210 | 0.108 | **VG-Flow** | 80 | 70.4 | 99229 | 0.670 |
| PredictHaplo | 8 | 53.3 | 10267 | 0.126 | PredictHaplo | 1 | 12.4 | - | 1.033 |
| ShoRAH | 493 | 26.3 | 10117 | 4.392 | ShoRAH | - | - | - | - |
| *6-strain Poliovirus* |  |  |  |  | *L = 200 kbp* |  |  |  |  |
| SAVAGE | 59 | 83.7 | 1643 | 0.019 | SAVAGE | 1910 | 66.0 | 975 | 0.016 |
| Virus-VG | 14 | 80.7 | 7428 | 0.064 | Virus-VG | - | - | - | - |
| **VG-Flow** | 12 | 90.2 | 7428 | 0.036 | **VG-Flow** | 167 | 64.1 | 201078 | 0.665 |
| PredictHaplo | 3 | 16.6 | - | 1.825 | PredictHaplo | - | - | - | - |
| ShoRAH | - | - | - | - | ShoRAH | - | - | - | - |

(a) Viral quasispecies benchmarks
(b) 8-strain mixtures of increasing genome length ( $L$ )

Compared to the state-of-the-art for full-length viral quasispecies reconstruction, we notice a clear advantage for VG-Flow in terms of target coverage and error rate. Table 1a shows that PredictHaplo and ShoRAH are unable to reconstruct all haplotypes in any of the simulated data sets. The strains that could be reconstructed have higher error rates than VG-Flow, as well as much higher frequency estimation errors. On the labmix data, shown in the Supplementary Material, PredictHaplo and ShoRAH both achieve a target coverage of 100%. In other words, they assemble each of the five HIV strains at full-length. However, PredictHaplo does so at almost twice the error rate of VG-Flow (1.066% for PredictHaplo versus 0.535% for VG-Flow) and the ShoRAH assembly has an even higher error rate of 3.910%. Moreover, ShoRAH greatly overestimates the number of strains in all data sets considered.

### Strain Abundance Estimation

Figure 3 shows the relative frequency estimation errors per method as a function of true abundance per strain, for each of the simulated data sets. Results are divided into bins (binsize=0.05) and average errors per bin^1^ are shown. Figure 3 highlights the advantage of de novo methods VG-Flow and Virus-VG, which have much smaller relative errors than PredictHaplo and ShoRAH. On the HCV and ZIKV data sets, VG-Flow and Virus-VG show nearly identical performance; on the Poliovirus data, we observe a small advantage for VG-Flow.

**Figure 3:**
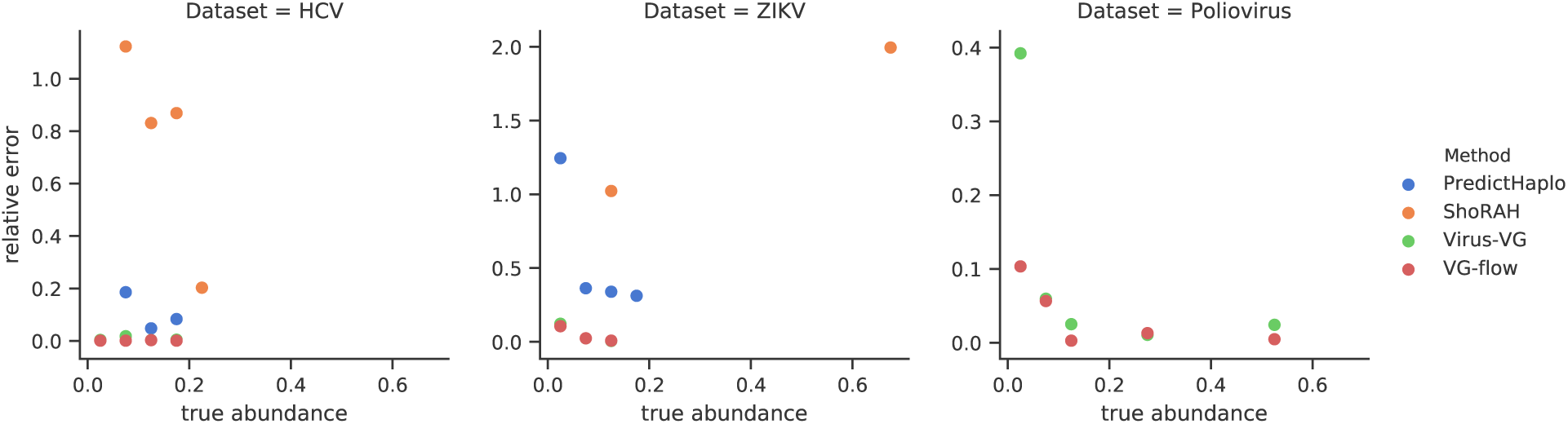
Abundance estimation results per data set. Relative errors are shown, calculated as |*x* − *x*\*|*/*(0.5(*x* + *x*\*)), where *x, x*\* represent the estimated and true values, respectively. Abundances were only evaluated for assemblies containing at least 2 full-length haplotypes and true abundance values were determined relative to the haplotypes present in the assembly.

### Long (Bacterial Sized) Genomes

In order to explore the limits of VG-Flow and to highlight its advantages over existing methods, we simulated 28 data sets with increasing genome sizes (2500 bp, 5000 bp, 10.000 bp, 20.000 bp, 40.000 bp, 100.000 bp, 200.000 bp) and an increasing number of strains (2, 4, 6, 8). Each data set has a total coverage of 1000x and haplotypes have a pairwise divergence of 1%. For data sets of 2, 4, 6, and 8 strains, the relative strain abundances were set to ratios of 1:2, 1:2:3:4, 1:2:3:4:5:6, and 1:2:3:4:5:6:7:8, respectively. Although the genome sequences are artificial, allowing us to vary genome size and number of strains flexibly, the relative abundances and pairwise divergence reflect plausible real-world, and challenging scenarios in metagenomics [11].

Table 1b presents assembly results for all methods on a selection of these data sets; as expected, none of Virus-VG, ShoRAH and PredictHaplo were able to process 8-strain mixtures in reasonable time for genomes ≥ 5 kbp, 20 kbp and 200 kbp, respectively. Only VG-Flow was able to assemble genomes beyond 10 kbp at sufficient target coverage, accurate NG50 and errors below the sequencing error rate (1%) which documents the quality of the strains. The overestimation of the number of strains can be attributed to ‘ambiguity islands’ between strains in varying genomic regions, and which can only be bridged using sufficiently long reads. Note also that the number of possible full-length paths overall amounts to roughly 8^10^, which puts VG-flow’s estimates into perspective, rendering them very reasonable. In the Supplementary Material we provide a more detailed runtime and scalability evaluation, as well as additional experiments.

### Practical runtime and scalability analysis

When Virus-VG was able to reconstruct full-length haplotypes, we measured a decrease in haplotype reconstruction time of 9.2–92% for VG-flow relative to Virus-VG. Thanks to the efficiency in the later steps, total runtime for VG-Flow is dominated by the contig variation graph construction step. Since these steps are shared by VG-Flow and Virus-VG, we observe near-identical runtimes overall (between 3.6–12.5 CPU hours and peak memory usage is between 0.6–0.9 GB on the viral quasispecies benchmarks), *for data sets which Virus-VG was able to process*. For the larger genomes (up to 200 kbp) VG-Flow used approximately 70 CPU hours, while Virus-VG could not finish those. Also, both approaches require pre-assembled contigs, for which we used SAVAGE (see the Supplement for results using pre-assembled contigs from SPAdes [17]). The generation of haplotype-aware contigs can be expensive (here: 30.6–276 CPU hours), see also ‘Remark on generation of haplotype-aware contigs’ in the Supplement, meaning that PredictHaplo is faster (2.0–7.4 CPU hours) overall (whereas ShoRAH is slower overall, 209–814 CPU hours) even when considering the generation of consensus reference genomes by VICUNA (0.07–0.44 CPU hours). Because all methods, except PredictHaplo, can be multithreaded, wall clock times are competitive in all these cases. Note that ShoRAH and PredictHaplo were unable to process genomes larger than 10 kbp and 100 kbp, respectively, within one week.

## Conclusions

While multiple approaches to reference-free viral quasispecies assembly have been introduced recently, efficient reconstruction of full-length haplotypes without using a reference genome is a major challenge. Although de novo methods have shown advantages over reference-guided tools, the resulting assemblies often consist of rather short contigs. In this paper, we have proposed VG-Flow as an efficient solution to extend pre-assembled contigs into full-length haplotypes, based on variation graphs as reference systems that allow for a bias-free consideration of all haplotypes involved. Benchmarking experiments have shown that VG-Flow outperforms the state-of-the-art in viral quasispecies reconstruction in terms of accuracy of haplotype sequences as well as abundance estimates. Moreover, we have shown that our method scales well to bacterial sized genomes, thus proving its potential for processing larger data sets like bacterial mixtures or metagenomic data. As soon as de novo assemblers such as [6, 17] are able to generate strain-specific contigs at this scale, VG-Flow can be applied to combine these contigs into full-length, strain-specific assemblies.

## Funding

This work was supported by the Netherlands Organisation for Scientific Research (NWO) through Vidi grant 679.072.309 and Gravitation Programme Networks 024.002.003.

## Footnotes

1 |*x* − *x*\*|*/*(0.5(*x* + *x*\*)), where *x, x*\* represent the estimated and true values, respectively

## References

[1] E. Domingo, J. Sheldon, and C. Perales. Viral quasispecies evolution. Microbiology and Molecular Biology Reviews, 76(2):159–216, Jun 2012.

[2] S. Crotty, C.E. Cameron, and R. Andino. RNA virus error catastrophe: direct molecular test by using ribavirin. Proceedings of the National Academy of Sciences, 98(12):6895–6900, 2001.

[3] M. Vignuzzi, J.K. Stone, J.J. Arnold, C.E. Cameron, and R. Andino. Quasispecies diversity determines pathogenesis through cooperative interactions in a viral population. Nature, 439:344–348, 2006.

[4] O. Zagordi, A. Bhattacharya, N. Eriksson, and N. Beerenwinkel. ShoRAH: estimating the genetic diversity of a mixed sample from next-generation sequencing data. BMC Bioinformatics, 12(1):119, 2011.

[5] A. Töpfer, O. Zagordi, S. Prabhakaran, V. Roth, E. Halperin, and N. Beerenwinkel. Probabilistic inference of viral quasispecies subject to recombination. Journal of Computational Biology, 20(2):113–123, 2013.

[6] J.A. Baaijens, A. Zine El Aabidine, E. Rivals, and A Schönhuth. De novo assembly of viral quasispecies using overlap graphs. Genome Research, 27(5):835–848, 2017.

[7] J.A. Baaijens, B. Van der Roest, J. Köster, L. Stougie, and A. Schönhuth. Full-length de novo viral quasispecies assembly through variation graph construction. Bioinformatics, 05 2019. btz443.

[8] J. Chen, Y. Zhao, and Y. Sun. De novo haplotype reconstruction in viral quasispecies using paired-end read guided path finding. Bioinformatics, 34(17):2927–2935, 2018.

[9] B. Paten, A.M. Novak, J.M. Eizenga, and E. Garrison. Genome graphs and the evolution of genome inference. Genome Research, 27(5):665–676, 2017.

[10] E. Garrison, J. Sirén, A.M Novak, G. Hickey, J.M. Eizenga, E.T. Dawson, W. Jones, S. Garg, C. Markello, M.F. Lin, B. Paten, and R. Durbin. Variation graph toolkit improves read mapping by representing genetic variation in the reference. Nature Biotechnology, 36:875–879, 2018.

[11] A. Sczyrba, Peter Hofmann, Peter Belmann, David Koslicki, Stefan Janssen, Johannes Dröge, Ivan Gregor, Stephan Majda, and A. McHardy. Critical assessment of metagenome interpretation - a benchmark of metagenomics software. Nature Methods, 14:1063–1071, 2017.

[12] S. Prabhakaran, M. Rey, O. Zagordi, N. Beerenwinkel, and V. Roth. HIV haplotype inference using a propagating dirichlet process mixture model. IEEE Transactions on Computational Biology and Bioinformatics, 11(1):182–191, 2014.

[13] M.C.F. Prosperi and M. Salemi. QuRe: software for viral quasispecies reconstruction from next-generation sequencing data. Bioinformatics, 28(1):132–133, Jan 2012.

[14] S. Ahn and H. Vikalo. aBayesQR: A bayesian method for reconstruction of viral populations characterized by low diversity. Journal of Computational Biology, 25(7):637–648, 2018.

[15] S. Barik, S. Das, and H. Vikalo. Qsdpr: Viral quasispecies reconstruction via correlation clustering. Genomics, 110(6):375–381, 2018.

[16] S. Knyazev, V. Tsyvina, A. Melnyk, A. Artyomenko, T. Malygina, Y.B. Porozov, E. Campbell, W.M. Switzer, P. Skums, and A. Zelikovsky. CliqueSNV: Scalable reconstruction of intra-host viral populations from NGS reads. bioRxiv:10.1101/264242, 2018.

[17] A. Bankevich, S. Nurk, D. Antipov, A.A. Gurevich, M. Dvorkin, A.S. Kulikov, V.M. Lesin, S.I. Nikolenko, S. Pham, A.D. Prijbelski, A.V. Pyshkin, A.V. Sirotkin, N. Vyahni, G. Tesler, P.A. Pevzner, and M.A. Alekseyev. SPAdes: A new genome assembly algorithm and its applications to single-cell sequencing. Journal of Computational Biology, 19(5):455–477, 2012.

[18] S. Nurk, D. Meleshko, A. Korobeynikov, and P.A. Pevzner. metaSPAdes: a new versatile metagenomic assembler. Genome Research, 27(5):824–834, 2017.

[19] S. Boisvert, F. Raymond, E. Godzaridis, F. Laviolette, and J. Corbeil. Ray meta: scalable de novo metagenome assembly and profiling. Genome Biology, 13(12):R122, 2012.

[20] Y. Peng, H.C. Leung, S.M. Yiu, and F.Y. Chin. Meta-IDBA: a de novo assembler for metagenomic data. Bioinformatics, 27(13):i94–i101, 2012.

[21] Dinghua Li, Chi-Man Liu, Ruibang Luo, Kunihiko Sadakane, and Tak-Wah Lam. MEGAHIT: an ultra-fast single-node solution for large and complex metagenomics assembly via succinct de Bruijn graph. Bioinformatics, 31(10):1674–1676, 01 2015.

[22] A.I. Tomescu, A. Kuosmanen, R. Rizzi, and V. Mäkinen. A novel min-cost flow method for estimating transcript expression with RNA-seq. BMC Bioinformatics, 14(5):S15, Apr 2013.

[23] R. Rizzi, A.I. Tomescu, and V. Mäkinen. On the complexity of minimum path cover with subpath constraints for multi-assembly. BMC Bioinformatics, 15(9):S5, 2014.

[24] E. Bernard, L. Jacob, J. Mairal, and J. Vert. Efficient RNA isoform identification and quantification from RNA-seq data with network flows. Bioinformatics, 30(17):2447–2455, 2014.

[25] M. Pertea, G.M. Pertea, C.M. Antonescu, T. Chang, J.T. Mendell, and S.L. Salzberg. StringTie enables improved reconstruction of a transcriptome from RNA-seq reads. Nature Biotechnology, 33:290–295, 2015.

[26] C. Trapnell, B.A. Williams, G. Pertea, A. Mortazavi, G. Kwan, M.J. van Baren, S.L. Salzberg, B.J. Wold, and L. Pachter. Transcript assembly and quantification by RNA-seq reveals unannotated transcripts and isoform switching during cell differentiation. Nature Biotechnology, 28:511–515, 2010.

[27] Y. Nesterov and A. Nemirovskii. Interior-point polynomial algorithms in convex programming, volume 13. SIAM, 1994.

[28] Ravindra K. Ahuja, Thomas L. Magnanti, and James B. Orlin. Network Flows: Theory, Algorithms, and Applications. Prentice-Hall, Inc., Upper Saddle River, NJ, USA, 1993.

[29] H. Li. Microbiome, metagenomics, and high-dimensional compositional data analysis. Annual Review of Statistics and Its Application, 2(1):73–94, 2015.

[30] Ana Conesa, Pedro Madrigal, Sonia Tarazona, David Gomez-Cabrero, Alejandra Cervera, Andrew McPherson, Michał Wojciech Szcześniak, Daniel J. Gaffney, Laura L. Elo, Xuegong Zhang, and Ali Mortazavi. A survey of best practices for RNA-seq data analysis. Genome Biology, 17:13, 2016.

[31] M.S. Lindner and B.Y. Renard. Metagenomic abundance estimation and diagnostic testing on species level. Nucleic Acids Research, 41(1):e10, 2012.

[32] M. Fischer, B. Strauch, and B.Y. Renard. Abundance estimation and differential testing on strain level in metagenomics data. Bioinformatics, 33(14):i124–i132, 2017.

[33] N.L. Bray, H. Pimentel, P. Melsted, and L. Pachter. Near-optimal probabilistic RNA-seq quantification. Nature Biotechnology, 34:525–527, 2016.

[34] G. Baier, E. Kohler, and M. Skutella. The k-splittable flow problem. Algorithmica, 42:231–248, 2005.

[35] B. Vatinlen, F. Chauvet, P. Chrétienne, and P. Mahey. Simple bounds and greedy algorithms for decomposing a flow into a minimal set of paths. European Journal of Operational Research, 185(3):1390–1401, 2008.

[36] M. Shao and C. Kingsford. Theory and a heuristic for the minimum path flow decomposition problem. IEEE/ACM Transactions on Computational Biology and Bioinformatics, PP(99):1–1, 2017.

[37] K. Kloster, P. Kuinke, M.P. O’Brien, F. Reidl, F. Sánchez Villaamil, B.D. Sullivan, and A. van der Poel. A practical fpt algorithm for flow decomposition and transcript assembly. CoRR, abs/1706.07851, 2017.

[38] William Cook and Paul Seymour. Tour merging via branch-decomposition. INFORMS Journal on Computing, 15(3):233–248, 2003.

[39] T. Bosman. A solution merging heuristic for the steiner problem in graphs using tree decompositions. In Evripidis Bampis, editor, *Experimental Algorithms*, pages 391–402, Cham, 2015. Springer International Publishing.

[40] C. Lee, C. Grasso, and M.F. Sharlow. Multiple sequence alignment using partial order graphs. Bioinformatics, 18(3):452–464, 2002.

[41] X. Yang, P. Charlebois, S. Gnerre, M. Coole, N. Lennon, J. Levin, J. Qu, E. Ryan, M. Zody, and M. Henn. De novo assembly of highly diverse viral populations. BMC Genomics, 13(1):475, 2012.

[42] John St. John. An illumina paired-end and mate-pair short read simulator. https://github.com/jstjohn/SimSeq, 2014.

[43] A. Gurevich, V. Saveliev, N. Vyahhi, and G. Tesler. QUAST: quality assessment tool for genome assemblies. Bioinformatics, 29(8):1072–1075, 2013.

