## Supplementary Material for "Strain-aware assembly of genomes from mixed samples using flow variation graphs"

Jasmijn A. Baaijens [1] and Leen Stougie [1,2] and Alexander Schönhuth [1,3,*]

[1]Centrum Wiskunde & Informatica, Amsterdam, Netherlands
[2]Vrije Universiteit, Amsterdam, Netherlands
[3]Utrecht University, Utrecht, Netherlands.

**4 SPAdes assemblies** **6**

**5 Contig abundance estimation** **8**

**6 Comparing greedy approaches** **15**

**7 Command lines** **20**

### 1 Further Related Work

The flow solution presents abundance estimates for the input contigs, which are of value in its own right in various mixed sample applications [1, 2, 3]. We use the contig abundance estimates in a combination of greedy algorithms to extract candidate haplotypes from the variation graph, where candidate haplotypes reflect concatenations of subpaths associated with the input contigs. Finally, we solve an optimization problem whose variables represent the haplotype abundances and the difference between read coverage and haplotype abundance is minimized over all nodes. Thus, we obtain a selection of candidate haplotypes that represents the quasispecies, along with haplotype abundance estimates.

Existing viral quasispecies assemblers include widely evaluated tools like [4, 5, 6], as well a variety of methods introduced more recently [7, 8, 9, 10, 11, 12]. These methods can be divided into two classes: reference-guided and reference-free (also referred to as *de novo*). De novo approaches do not require any prior information, such as a reference genome or knowledge of the quasispecies composition. This has been shown to have advantages over reference-guided reconstruction, since using a reference genome can induce significant biases. Especially at the time of a viral disease outbreak, an appropriate reference genome may not be available due to high mutation rates. However, most of the aforelisted tools are reference-guided; only [8], [9], and [11] present de novo approaches.

Moreover, many of these specialized viral quasispecies assemblers aim at single gene reconstruction, rather than whole genome assembly. In [9] we took a first step towards full-length de novo viral quasispecies assembly. There, we have shown that de novo haplotype reconstruction with integrated haplotype abundance estimation yields assemblies that are more complete, more accurate, and provide better abundance estimates. While this approach is guaranteed to find a selection of haplotypes that is optimal in terms of being compatible with the read coverages, its runtime is exponential in the number of contigs. Since the number of contigs generally increases on increasing genome length, we found [9] unsuitable for genomes larger than $\sim$ 10 kb. With VG-flow, we provide a reference-free solution to the full-length viral quasispecies reconstruction problem that scales well to longer genomes. As another benefit of the theoretical rigorosity of our problem formulation and efficiency of the solution, we also experience considerable improvements in terms of accuracy compared to existing tools.

Some of the challenges that have to be dealt with in viral quasispecies assembly can also be found in RNA transcipt assembly, where the goal is to reconstruct an unknown number of transcripts and predict the relative transcript abundances. Not surprisingly, many RNA transcipt assemblers define graph optimization problems similar to our flow formulation [13, 14, 15, 16, 17]. Although dealing with related problems, these methods cannot be applied in a viral quasispecies setting so easily: they require a collection of reference genomes representing all possible haplotypes as input, which is not available in our setting. Nevertheless, the theory behind these approaches is related to what we do. In [13], node and edge abundance errors are used to define a min-cost flow problem; note that this formulation does not take subpath constraints into account. On the other hand, [14] describes how subpath constraints can be incorporated into a minimum path cover formulation. This results in an optimization problem that is solvable in polynomial time, but does not minimize node abundance errors. We use the best of both worlds by defining an optimization problem that takes subpath constraints, minimizes node abundance errors, and is polynomially solvable; this establishes a theoretical novelty. Because this novelty gives way to different types of analyses in other settings, and immediately connects to extensively treated theoretical issues [13, 14], we feel that it is of value also in its own right.

Among more generic assemblers, [18] has been shown to be capable of reconstructing individual haplotypes from mixed samples, up to a certain degree. This method was designed for bacterial

| Data set | Data type | Virus type | Genome size | Strain count | Strain abundance | Pairwise divergence |
|---|---|---|---|---|---|---|
| HCV mix | Simulated | HCV-1a | 9273–9311 bp | 10 | 5–19% | 6–9% |
| ZIKV mix | Simulated | ZIKV | 10251–10269 bp | 15 | 2–13% | 1–10% |
| Poliovirus mix | Simulated | Poliovirus | 7428–7460 bp | 6 | 1.6–51% | 1.2–7% |
| Labmix | Real | HIV-1 | 9478–9719 bp | 5 | 10–30% | 1–6% |

Table 1: Quasispecies characteristics of benchmarking data sets. All data sets consist of Illumina Miseq reads with an average sequencing depth of 20.000x.

genomes and scales well to human genomes, but is unable to reconstruct low-frequent haplotypes [8]. Haplotype-aware assembly of metagenomes is a big challenge, which tends to result in scattered genome fragments and missing strains [19]. Metagenomic assemblers such as [20, 21, 22, 23] aim to reconstruct mixtures of viral and bacterial populations at strain level. The contigs obtained with these methods, or any other assembler, can also be used as input for VG-flow. Although we emphasize the reconstruction of viral quasispecies, the mathematical framework presented here is generic and could be applied in other scenarios as well; as such, VG-flow has the potential to make a big step ahead in haplotype-aware genome assembly in general.

### 2 Data sets

We evaluate performance of VG-flow on three simulated viral quasispecies data sets from [9] and one real HIV benchmark presented in [24], also referred to as the *labmix*. The simulated data sets are based on true genomic sequences from the NCBI nucleotide database; the characteristics of all data sets (virus type, genome size, number of strains, relative strain abundances, and pairwise divergence) are described in Table 1. All data sets consist of Illumina Miseq reads with an average sequencing depth of 20.000x. For each data set, including the labmix, the true haplotypes and their relative abundances are known.

### 3 Extended results

We evaluate all assemblies by comparing the assembled contigs to the ground truth sequences using QUAST [25]. This assembly evaluation tool aligns the assembled contigs to the true haplotypes, which are provided as a reference, and calculates several standard evaluation metrics. For each assembly, we report the number of contigs, percent target genomes covered, N50, NG50, and error rate. If an assembly consists of only full-length contigs, the number of contigs can be interpreted as the estimated number of strains. Target genome coverage is defined as the percentage of aligned bases in the true haplotypes, where a base is considered aligned if there is at least one contig with at least one alignment to this base. The N50 and NG50 measures reflect assembly contiguity. N50 is defined as the length for which all contigs in the assembly of at least this length together add up to at least half of the total assembly size. NG50 is calculated in a similar fashion, except that the sum of contig lengths is required to cover at least half of the total target length. Error rates are equal to the sum of mismatch rate, indel rate, and N-rate (ambiguous bases). We do not report unaligned bases or misassemblies as we did not encounter any of these.

In addition to the above QUAST assembly metrics, we evaluate strain abundance estimates by comparing estimated values to true strain abundances. For each assembly, let $n$ be the number of true strains and let $x_i, x'_i$ denote the estimated and true abundance, respectively, of strain $i$.

For each ground truth haplotype, the abundance estimates of sequences assigned to this haplotype were summed to obtain the strain abundance estimate $x_i$. We only evaluate abundances for strains that are present in the assembly (i.e. $x_i > 0$) since we cannot expect an assembler to estimate the abundance of a missing strain. Therefore, the true strain abundance values $x'_i$ are also normalized, taking only the assembled sequences into account. Then, we calculate the absolute frequency error (AFE) and the relative frequency error (RFE) a follows:

$$\begin{aligned} \text{AFE} &= \sum_{i \in I} \frac{|x_i - x'_i|}{|I|}, \\ \text{RFE} &= \sum_{i \in I} \frac{|x_i - x'_i|}{|I| \cdot x'_i}, \text{ where} \\ I &= \{i \in [n] : x_i > 0\} \end{aligned}$$

##### 3.1 Simulated data

The frequency estimation errors (AFE and RFE) in Table 2 show that the increase in assembly accuracy also leads to lower frequency estimation errors.

##### 3.2 Real data

On real data (Table 3) we observe that VG-flow produces the same number of contigs as Virus-VG, leading to identical target coverage and N50 values; the only difference between the assemblies is a slightly lower NG50 for VG-flow (4608 versus 4642) and a slightly higher error rate (0.535% versus 0.324%). These differences may be explained by the highly uneven coverage of this data set, which affects the contig abundance estimation and hence also the greedy path extraction. However, the contigs produced by VG-flow are much longer than the input contigs, with the N50 value more than doubled.

##### 3.3 Runtime and scalability

Table 4 presents runtime and memory usage for SAVAGE, Virus-VG, VG-flow, PredictHaplo, and ShoRAH on viral quasispecies benchmarks. Figure 1 compares runtimes for the haplotype reconstruction process of VG-flow and Virus-VG (excluding de novo assembly and contig-variation graph construction, which are shared by the two methods). Figure 2 shows total runtimes for VG-flow (including de novo assembly with SAVAGE) as a function of genome size for data sets with an increasing number of strains. Note that for assembling the 40.000, 100.000, and 200.000 bp genomes we had to use the SAVAGE `--split` parameter to split the data over 2, 5, and 10 patches, respectively. Increasing the number of patches further would lead to too little coverage per patch, hence, for larger genomes, a different assembler is required.

*Remark on generation of haplotype-aware contigs.* Currently, the limiting factor for processing genomes larger than 200.000 bp with VG-Flow is the pre-assembly step. VG-Flow requires pre-assembled strain-specific contigs as input and we use SAVAGE [8] for this. SAVAGE has proven to produce assemblies of very high quality, but this assembler does not scale well to large genomes. Inspired by results from [8], we experimented with SPAdes [18] assemblies as input for VG-Flow. Although SPAdes does not produce strain-specific contigs as well as SAVAGE, it performs reasonably well and VG-Flow is able to build full-length haplotypes from these contigs (see 'SPAdes assemblies' further below).

| | # contigs* | target (%) | N50 | NG50 | ER(%) | AFE(%) | RFE(%) |
|---|---|---|---|---|---|---|---|
| SAVAGE | 26 | 99.4 | 8964 | 8964 | 0.001 | - | - |
| Virus-VG | 10 | 99.3 | 9281 | 9203 | 0.001 | 0.1 | 0.9 |
| VG-flow | 10 | 99.3 | 9281 | 9203 | 0.001 | 0.0 | 0.2 |
| PredictHaplo | 9 | 73.8 | 7636 | 7608 | 0.059 | 0.9 | 11.3 |
| ShoRAH | 639 | 56.9 | 7570 | 7570 | 4.294 | 8.5 | 64 |

(a) 10-strain HCV mixture

| | # contigs* | target (%) | N50 | NG50 | ER(%) | AFE(%) | RFE(%) |
|---|---|---|---|---|---|---|---|
| SAVAGE | 100 | 98.8 | 2954 | 3801 | 0.023 | - | - |
| Virus-VG | 20 | 92.8 | 10202 | 10210 | 0.115 | 0.3 | 6.0 |
| VG-flow | 21 | 92.8 | 10193 | 10210 | 0.108 | 0.3 | 5.4 |
| PredictHaplo | 8 | 53.3 | 10270 | 10267 | 0.126 | 4.9 | 69 |
| ShoRAH | 493 | 26.3 | 10117 | 10117 | 4.392 | 39 | 229 |

(b) 15-strain ZIKV mixture

| | # contigs* | target (%) | N50 | NG50 | ER(%) | AFE(%) | RFE(%) |
|---|---|---|---|---|---|---|---|
| SAVAGE | 59 | 83.7 | 1089 | 1643 | 0.019 | - | - |
| Virus-VG | 14 | 80.7 | 7316 | 7428 | 0.064 | 0.6 | 12.8 |
| VG-flow | 12 | 90.2 | 7316 | 7428 | 0.036 | 0.3 | 3.5 |
| PredictHaplo | 3 | 16.6 | 7461 | - | 1.825 | - | - |

(c) 6-strain Poliovirus mixture

Table 2: Assembly results on simulated data (Illumina MiSeq, 20.000x coverage). ER = Error Rate (N's + mismatches + indels), AFE = Absolute Frequency Error, RFE = Relative Frequency Error. Frequency errors were only computed for assemblies containing at least 2 full-length haplotypes. *If contigs are full-length, this number reflects the estimated number of strains in the quasispecies.

### 4 SPAdes assemblies

Table 5 presents statistics for assemblies obtained with SPAdes (in `--careful` mode) and the haplotypes generated with VG-flow using the SPAdes assemblies as input.

| | # contigs* | target (%) | N50 | NG50 | ER(%) |
|---|---|---|---|---|---|
| SAVAGE | 68 | 97.9 | 1026 | 1450 | 0.066 |
| Virus-VG | 23 | 90.6 | 2130 | 4642 | 0.324 |
| VG-flow | 23 | 90.6 | 2130 | 4608 | 0.535 |
| PredictHaplo | 6 | 100.0 | 8825 | 8825 | 1.066 |
| ShoRAH | 250 | 100.0 | 8775 | 8775 | 3.910 |

Table 3: Assembly results on the labmix (5-strain HIV mixture, real Illumina MiSeq, 20.000x coverage).

| | HCV | | ZIKV | | Poliovirus | | Labmix | |
|---|---|---|---|---|---|---|---|---|
| | CPU time (hours) | Peak mem (GB) | CPU time (hours) | Peak mem (GB) | CPU time (hours) | Peak mem (GB) | CPU time (hours) | Peak mem (GB) |
| SAVAGE | 38.6 | 26 | 30.6 | 13 | 60.2 | 3.3 | 276 | 7.2 |
| Virus-VG | 5.9 | 0.9 | 12.5 | 0.8 | 3.6 | 0.6 | 10 | 0.6 |
| VG-flow | 5.9 | 0.9 | 12.5 | 0.8 | 3.6 | 0.6 | 10 | 0.6 |
| PredictHaplo | 2.7 | 1.1 | 7.4 | 1.1 | 2.0 | 0.8 | 4.9 | 1.0 |
| ShoRAH | 209 | 8.9 | 814 | 10 | - | - | 351 | 10.7 |

Table 4: Runtime and memory usage on viral quasispecies benchmarks. Note that VG-flow and Virus-VG runtimes are completely determined by the contig-variation graph construction step.

| | # contigs* | target (%) | N50 | NG50 | ER(%) | AFE(%) | RFE(%) |
|---|---|---|---|---|---|---|---|
| SPAdes | 27 | 94.3 | 8862 | 8397 | 0.008 | - | - |
| SPAdes + VG-flow | 10 | 98.0 | 9311 | 9311 | 0.080 | 0.0 | 0.4 |

(a) 10-strain HCV mixture

| | # contigs* | target (%) | N50 | NG50 | ER(%) | AFE(%) | RFE(%) |
|---|---|---|---|---|---|---|---|
| SPAdes | 259 | 59.7 | 216 | 252 | 0.195 | - | - |
| SPAdes + VG-flow | 18 | 52.2 | 3018 | 1337 | 0.373 | 9.0 | 36.0 |

(b) 6-strain Poliovirus mixture

| | # contigs* | target (%) | N50 | NG50 | ER(%) | AFE(%) | RFE(%) |
|---|---|---|---|---|---|---|---|
| $L = 2500$ | | | | | | | |
| SPAdes | 5 | 76.0 | 930 | 778 | 0.053 | - | - |
| SPAdes + VG-flow | 3 | 100 | 2497 | 2497 | 0.220 | 0.7 | 1.6 |
| $L = 5000$ | | | | | | | |
| SPAdes | 9 | 67.0 | 1892 | 695 | 0.015 | - | - |
| SPAdes + VG-flow | 3 | 50.0 | 4404 | 4404 | 0 | 0.0[a] | 0.0 |
| $L = 10.000$ | | | | | | | |
| SPAdes | 27 | 81.3 | 696 | 678 | 0.025 | - | - |
| SPAdes + VG-flow | 4 | 100 | 9997 | 10005 | 0.180 | 0.6 | 1.9 |
| $L = 20.000$ | | | | | | | |
| SPAdes | 42 | 68.0 | 1231 | 854 | 0.055 | - | - |
| SPAdes + VG-flow | 5 | 100 | 20002 | 20014 | 0.240 | 0.7 | 1.7 |
| $L = 40.000$ | | | | | | | |
| SPAdes | 94 | 82.9 | 952 | 845 | 0.008 | - | - |
| SPAdes + VG-flow | 11 | 55.0 | 20919 | 20924 | 0.259 | 0.0[a] | 0.0[a] |
| $L = 100.000$ | | | | | | | |
| SPAdes | 248 | 80.5 | 1031 | 890 | 0.055 | - | - |
| SPAdes + VG-flow | 20 | 73.1 | 46263 | 58056 | 0.321 + 1ME[b] | 0.0[a] | 0.0[a] |
| $L = 200.000$ | | | | | | | |
| SPAdes | 508 | 81.9 | 898 | 797 | 0.039 | - | - |
| SPAdes + VG-flow | 36 | 52.9 | 60475 | 150143 | 0.231 + 16ME[b] | 0.0[a] | 0.0[a] |

(c) Simulated mixtures of 2 strains at 1% divergence with genome size $L$

[a]Only 1 strain reconstructed in full-length, resulting in perfect frequency estimation

[b]ME: misassembly events; a position where the left and right flanking sequences align to the true genomes with a gap or overlap of more than 1 kbp, or align to different strands, or even align to different strains.

Table 5: Assembly results using SPAdes contigs as input for VG-flow. Contig-variation graph construction from SPAdes assemblies did not succeed for the simulated ZIKV data set and the labmix (real HIV data). ER = Error Rate (N's + mismatches + indels), AFE = Absolute Frequency Error, RFE = Relative Frequency Error. *If contigs are full-length, this number reflects the estimated number of strains in the quasispecies.

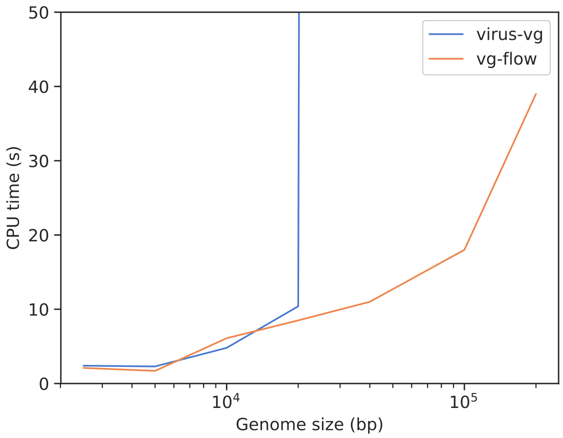

Figure 1: VG-flow and Virus-VG optimization runtimes on data sets of increasing genome size (2500, 5000, 10.000, 20.000, 40.000, 100.000, 200.000) containing 2 strains. The runtimes shown here exclude contig-variation graph construction times and SAVAGE assembly time (these steps are equally applicable to VG-flow and Virus-VG). At a genome size of 40.000 bp the runtime of Virus-VG increased to >1h.

### 5 Contig abundance estimation

We estimate contig abundances by solving a flow-like optimization problem: variables represent flow values on the edges of the flow graph and we impose flow constraints, while the objective function evaluates the difference between estimated contig abundances and read coverage for every node in the variation graph. Lemma 5.1 shows that the objective function is convex, hence the optimization problem is polynomial time solvable.

**Lemma 5.1.** *Let $E = \{e_1, \ldots, e_m\}$ and let $x = (x_{e_1}, \ldots, x_{e_m}) \in \mathbb{R}^m$. The function*

$$f(x) := \sum_{u \in U} |a_u - \sum_{\{e \in E | u \in e\}} d_e x_e|$$

*is convex in the flow variables $x_e$, $e \in E$.*

*Proof.* To prove convexity, we need to show that $f(\lambda x + (1-\lambda)x') \leq \lambda f(x) + (1-\lambda) f(x')$ for any $x, x' \in \mathbb{R}^n$ and $\lambda \in [0,1]$. Hence, it is sufficient to show that, for any $u \in U$ and $\lambda \in [0,1]$,

$$\left| \lambda \left( a_u - \sum_{\{e \in E | u \in e\}} d_e x_e \right) + (1-\lambda) \left( a_u - \sum_{\{e \in E | u \in e\}} d_e x'_e \right) \right| \leq \lambda \left| a_u - \sum_{\{e \in E | u \in e\}} d_e x_e \right| + (1-\lambda) \left| a_u - \sum_{\{e \in E | u \in e\}} d_e x'_e \right|$$

which follows immediately from the triangle inequality. □

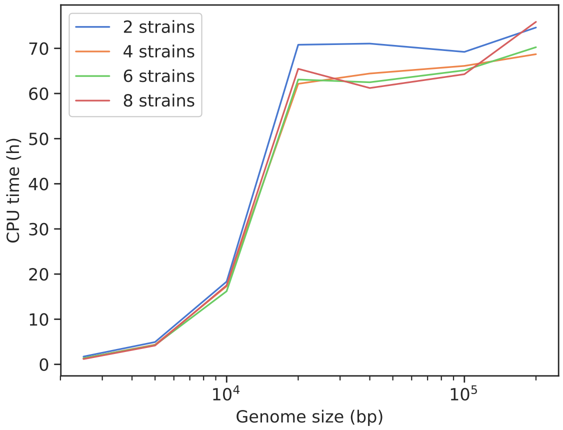

Figure 2: Total runtimes (SAVAGE + contig-variation graph construction + VG-flow optimization) on data sets of increasing genome size (2500, 5000, 10.000, 20.000, 40.000, 100.000, 200.000) and number of strains (2, 4, 6, 8). Note that total runtime is mainly determined by SAVAGE runtime (78–99%) and increases linearly with the genome size (the x-axis is plotted on a logarithmic scale). With a smaller number of strains, the coverage per strain is higher and the overlap graph used in SAVAGE becomes more dense. This leads to a higher runtime for SAVAGE as the number of strains decreases. For genome sizes 40.000, 100.000, and 200.000 the data set had to be split into 2, 5, and 10 patches, respectively, to run SAVAGE. This leads to lower coverage per strain, hence relatively faster processing by SAVAGE. Since SAVAGE runtime is the main component of total runtime, we also see that total runtime follows a different curve from 40.000 bp onwards.

Although the main goal of the contig abundance computation is to enable full-length haplotype reconstruction, the contig abundances are also of interest for metagenomic quantification purposes. We evaluate the estimated contig abundances by computing a "true abundance" for each contig in our simulated benchmarks. Even though the true haplotypes are known, computing true abundances is quite complicated: some contigs span conserved regions and hence map to multiple haplotypes, while for other regions we observe multiple contigs mapping to the same region due to sequencing or assembly errors. The most difficult case occurs when a contig maps equally well to multiple haplotypes. We could not find a way to properly compute the true abundance of such a contig, hence we only evaluate abundances for error-free contigs. Fortunately, this still constitutes a major part of the assemblies (92% for HCV, 78% for ZIKV, 89% for Poliovirus). The only ambiguity we have to deal with is contigs mapping without errors to multiple haplotypes, in which case we sum the abundances of the respective haplotypes to obtain the true contig abundance.

Since the computed contig abundances are absolute numbers (e.g. 536x), we represent the true abundances as absolute numbers as well. For a given contig, let $x_1$ be the estimated abundance and let $x_2$ be the true abundance, then we compute the relative error $|x_1 - x_2|/(0.5(x_1 + x_2))$.

Figures 3,4 and 5 show the relative errors per data set as a function of true abundance, estimated abundance, and contig alignment position, respectively. Figures 6 and 8 show the distribution of errors for all simulated data sets. Figures 7 and 9 present the distribution of errors when only contigs with an estimated abundance of at least 1% of the total sequencing depth are taken into account. Note that by default, any haplotypes with an estimated abundance below 1% are removed from the reconstructed haplotypes by VG-flow.

We observe higher error rates as data sets become more complex (more strains, low abundance, low pairwise divergence). Especially when assemblies are incomplete (i.e. $\ll 100\%$ target coverage) error rates increase. A likely explanation is that, on the one hand, contig abundance estimation becomes harder, while on the other hand, the true abundances become less reliable. When haplotypes are not fully reconstructed, reads belonging to these haplotypes will align to nodes of the variation graph corresponding to contigs belonging to other haplotypes. This has a great impact on abundance estimation. Since the majority of contig abundance estimates is very close to the true abundance, the erroneous estimates do not hamper the haplotype reconstruction process.

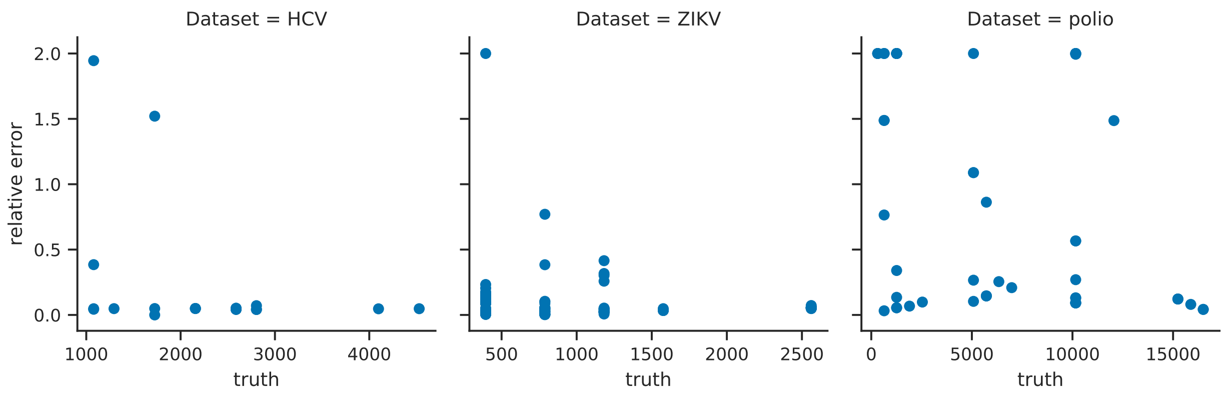

Figure 3: Relative contig abundance estimation errors versus true abundance

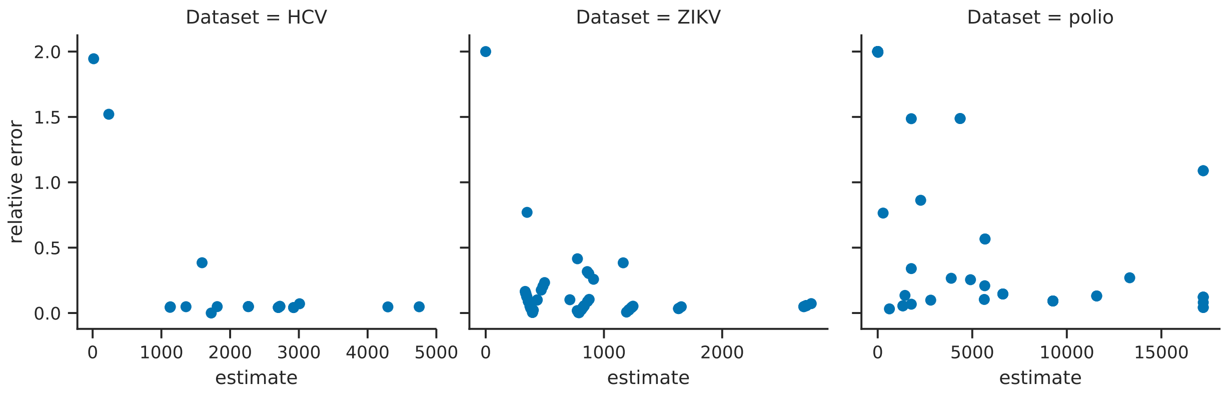

Figure 4: Relative contig abundance estimation errors versus estimated abundance

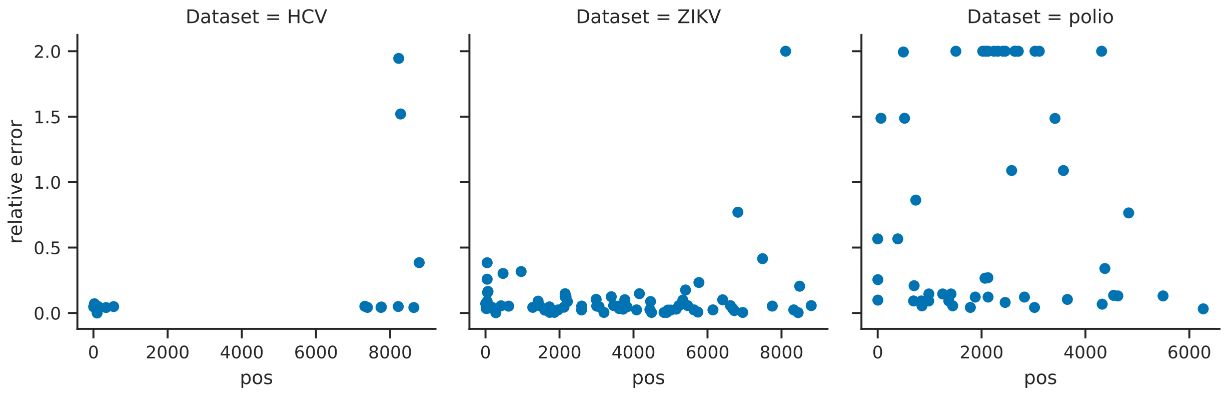

Figure 5: Relative contig abundance estimation errors versus contig alignment position

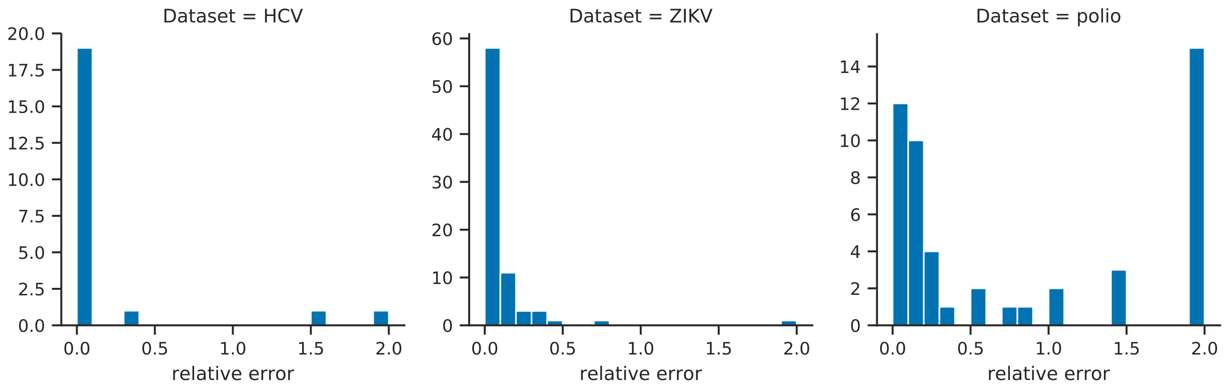

Figure 6: Distribution of relative contig abundance estimation errors

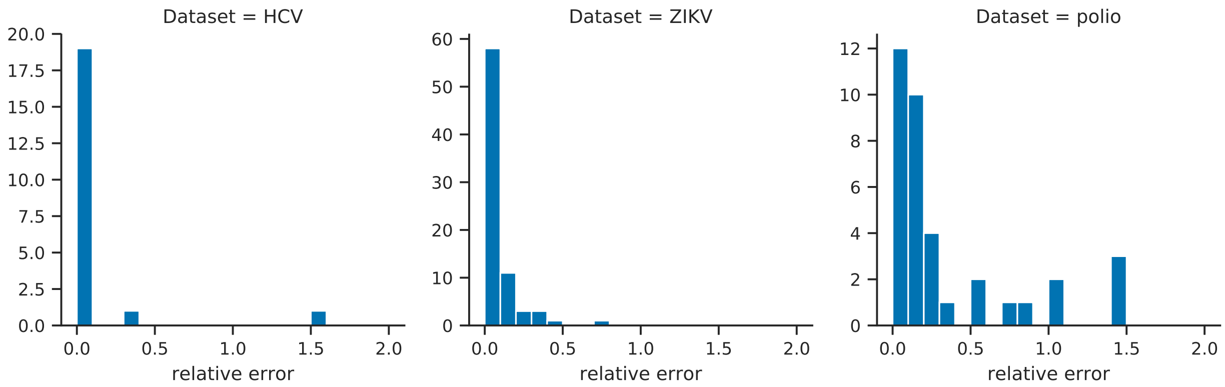

Figure 7: Distribution of relative contig abundance estimation errors for contigs with estimated abundance greater than 1%.

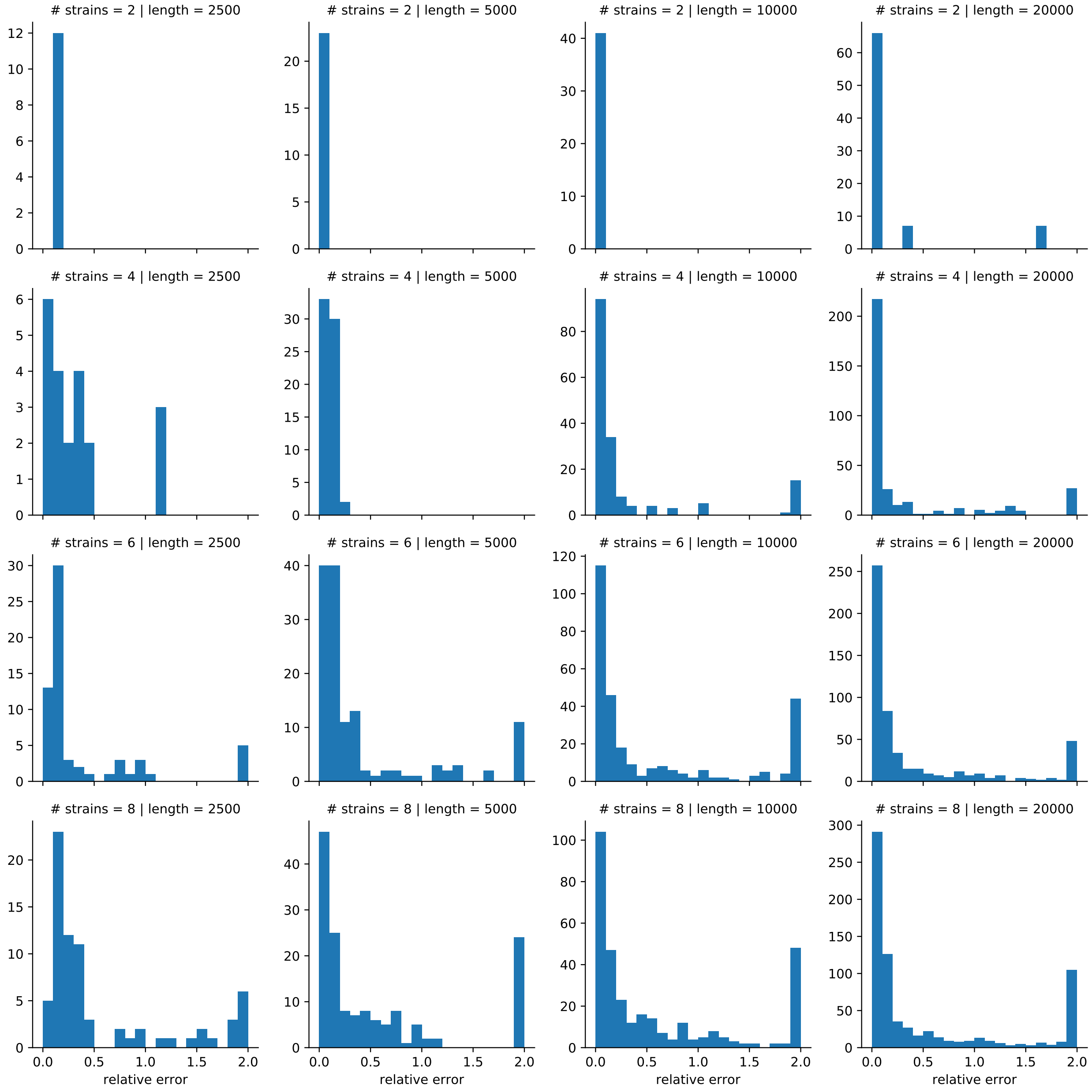

Figure 8: Distribution of relative contig abundance estimation errors

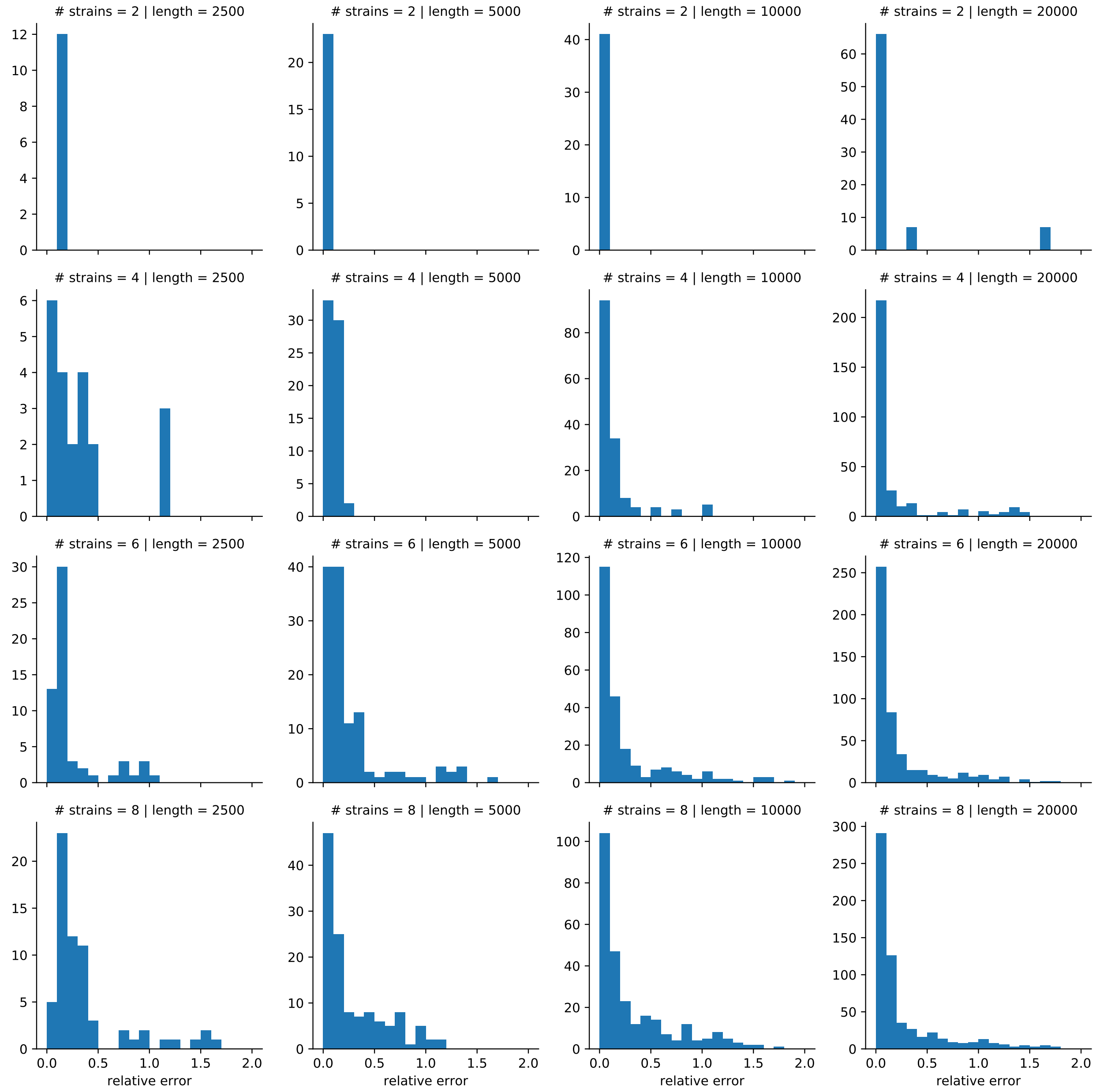

Figure 9: Distribution of relative contig abundance estimation errors for contigs with estimated abundance greater than 1%.

### 6 Comparing greedy approaches

Tables 6 and 7 show results for all greedy approaches on the simulated and real quasispecies benchmarks, respectively. Barplots in Figure 10 present results on the simulated data sets of increasing genome size and number of strains, for each of the different haplotype reconstruction approaches (Virus-VG, GreedyMaxCapacity, GreedyMinCapacity, GreedyShortestPaths, GreedyALL). Contig variation graph statistics corresponding to the data sets with a genome size of 5000 bp, as well as graph construction times and haplotype reconstruction times are presented in Table 8.

| | # contigs* | target (%) | N50 | NG50 | ER(%) | AFE(%) | RFE(%) |
|---|---|---|---|---|---|---|---|
| SAVAGE | 26 | 99.4 | 8964 | 8964 | 0.001 | - | - |
| Virus-VG | 10 | 99.3 | 9281 | 9203 | 0.001 | 0.1 | 0.9 |
| PredictHaplo | 9 | 73.8 | 7636 | 7608 | 0.059 | 0.9 | 11.3 |
| ShoRAH | 639 | 56.9 | 7570 | 7570 | 4.294 | 8.5 | 64 |
| GreedyMaxCapacity | 11 | 99.4 | 9281 | 9269 | 0.025 | 0.0 | 0.2 |
| GreedyMinCapacity | 11 | 99.3 | 9204 | 9203 | 0.001 | 0.0 | 0.2 |
| GreedyShortestPaths | 10 | 99.3 | 9281 | 9203 | 0.005 | 0.0 | 0.2 |
| GreedyALL | 10 | 99.3 | 9281 | 9203 | 0.001 | 0.0 | 0.2 |

(a) 10-strain HCV mixture

| | # contigs* | target (%) | N50 | NG50 | ER(%) | AFE(%) | RFE(%) |
|---|---|---|---|---|---|---|---|
| SAVAGE | 100 | 98.8 | 2954 | 3801 | 0.023 | - | - |
| Virus-VG | 20 | 92.8 | 10202 | 10210 | 0.115 | 0.3 | 6.0 |
| PredictHaplo | 8 | 53.3 | 10270 | 10267 | 0.126 | 4.9 | 69 |
| ShoRAH | 493 | 26.3 | 10117 | 10117 | 4.392 | 39 | 229 |
| GreedyMaxCapacity | 17 | 86.2 | 10202 | 10202 | 0.036 | 0.4 | 7.8 |
| GreedyMinCapacity | 18 | 86.2 | 10202 | 10202 | 0.043 | 0.4 | 6.7 |
| GreedyShortestPaths | 21 | 92.8 | 10193 | 10210 | 0.118 | 0.3 | 5.5 |
| GreedyALL | 21 | 92.8 | 10193 | 10210 | 0.108 | 0.3 | 5.4 |

(b) 15-strain ZIKV mixture

| | # contigs* | target (%) | N50 | NG50 | ER(%) | AFE(%) | RFE(%) |
|---|---|---|---|---|---|---|---|
| SAVAGE | 59 | 83.7 | 1089 | 1643 | 0.019 | - | - |
| Virus-VG | 14 | 80.7 | 7316 | 7428 | 0.064 | 0.6 | 12.8 |
| PredictHaplo | 3 | 16.6 | 7461 | - | 1.825 | - | - |
| GreedyMaxCapacity | 8 | 73.5 | 7390 | 7390 | 0.027 | 1.5 | 14 |
| GreedyMinCapacity | 11 | 89.8 | 7256 | 7390 | 0.041 | 0.3 | 3.8 |
| GreedyShortestPaths | 12 | 88.5 | 7376 | 7390 | 0.059 | 0.3 | 5.3 |
| GreedyALL | 12 | 90.2 | 7316 | 7428 | 0.036 | 0.3 | 3.5 |

(c) 6-strain Poliovirus mixture

Table 6: Assembly results on simulated data (Illumina MiSeq, 20.000x coverage). ER = Error Rate (N's + mismatches + indels), AFE = Absolute Frequency Error, RFE = Relative Frequency Error. *If contigs are full-length, this number reflects the estimated number of strains in the quasispecies.

| | # contigs* | target (%) | N50 | NG50 | ER(%) |
|---|---|---|---|---|---|
| SAVAGE | 68 | 97.9 | 1026 | 1450 | 0.066 |
| Virus-VG | 23 | 90.6 | 2130 | 4642 | 0.324 |
| PredictHaplo | 6 | 100.0 | 8825 | 8825 | 1.066 |
| ShoRAH | 250 | 100.0 | 8775 | 8775 | 3.910 |
| GreedyMaxCapacity | 21 | 90.7 | 2336 | 4642 | 0.274 |
| GreedyMinCapacity | 23 | 90.6 | 2130 | 4608 | 0.885 |
| GreedyShortestPaths | 23 | 90.6 | 2130 | 4608 | 0.896 |
| GreedyALL | 23 | 90.6 | 2130 | 4608 | 0.535 |

Table 7: Assembly results on the labmix (5-strain HIV mixture, real Illumina MiSeq, 20.000x coverage).

| | 2 strains | 4 strains | 6 strains | 8 strains |
|---|---|---|---|---|
| *Graph statistics* | | | | |
| # nodes | 141 | 274 | 413 | 537 |
| # edges | 188 | 367 | 555 | 723 |
| # candidate paths | 6 | 308 | - | - |
| *Graph construction time* | 149 | 383 | 919 | 867 |
| *Haplotype reconstruction time* | | | | |
| Virus-VG | 2.3 | 12 | - | - |
| GreedyMaxCapacity | 2.1 | 8.8 | 13 | 15 |
| GreedyMinCapacity | 3.0 | 9.6 | 13 | 18 |
| GreedyShortestPaths | 1.9 | 8.8 | 14 | 18 |
| GreedyALL | 2.9 | 7.8 | 15 | 21 |

Table 8: Graph statistics, graph construction and haplotype reconstruction runtimes (CPU seconds) for data sets with an increasing number of strains (genome size = 5000 bp).

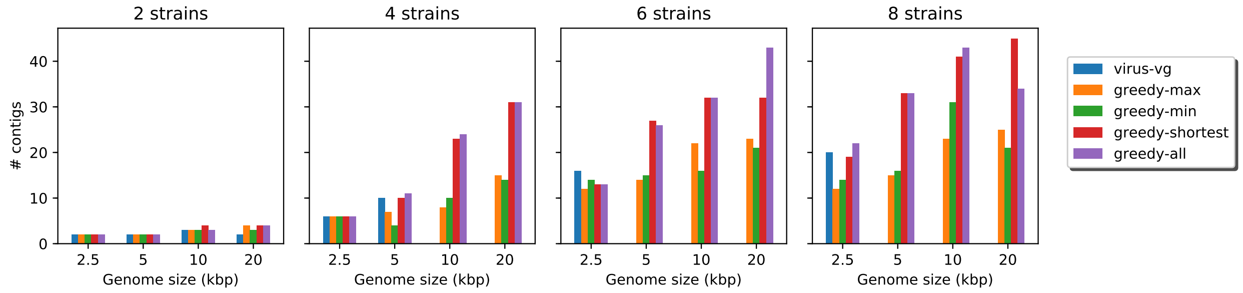

(a) # contigs

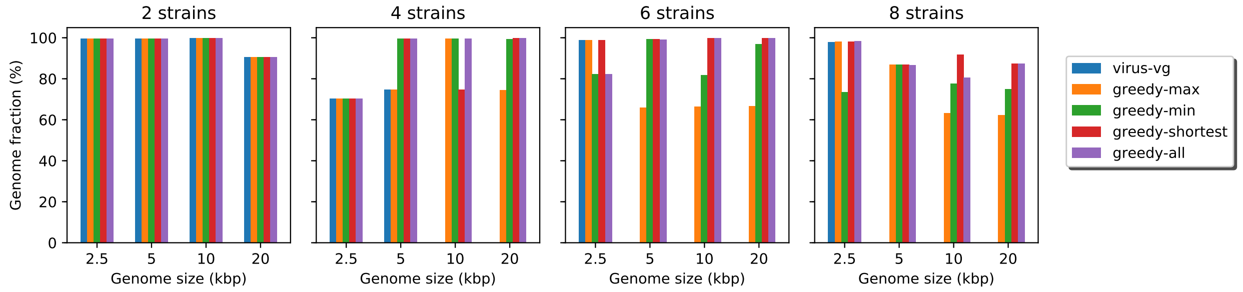

(b) Genome fraction (%)

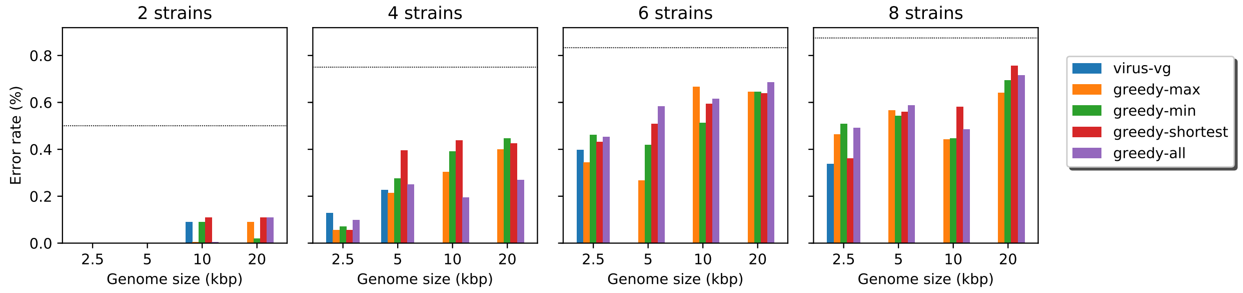

(c) Error rate (%) – *Dashed lines indicate expected error rates when assigning variants randomly.*

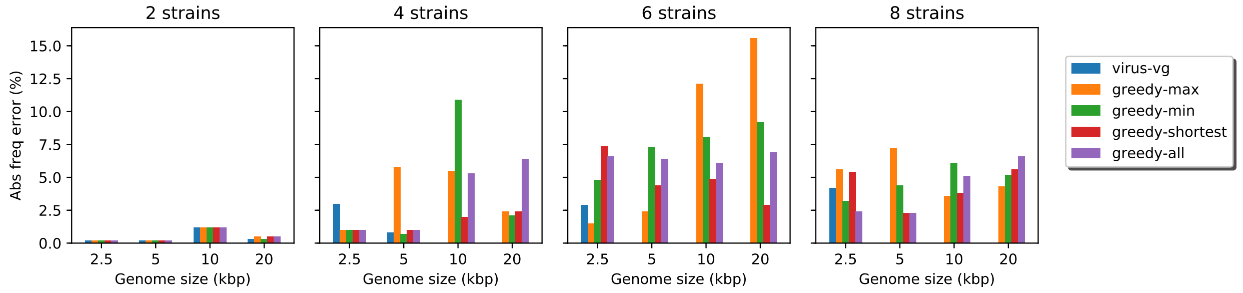

(d) Absolute frequency error rate (%)

Figure 10: Assembly statistics for data sets of increasing genome size (2500 bp, 5000 bp, 10.000 bp, 20.000 bp) and increasing number of strains (2, 4, 6, 8). Virus-VG results are missing for samples of 6 and 8 strains for genomes of size 5000 bp and up, as well as for samples of 4 strains for genomes of size at least 10.000 bp, because path enumeration became infeasible.

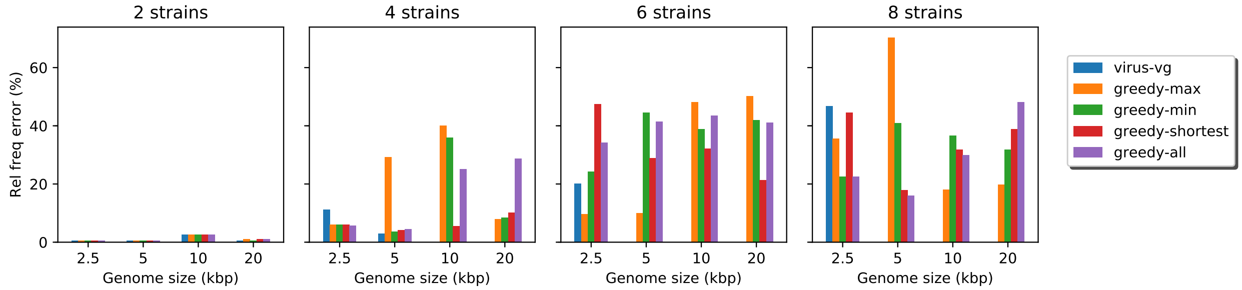

(e) Relative frequency error rate (%)

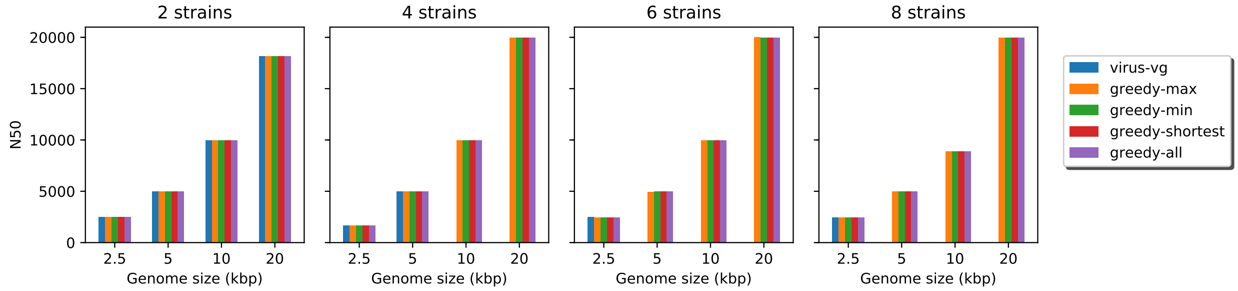

(f) N50

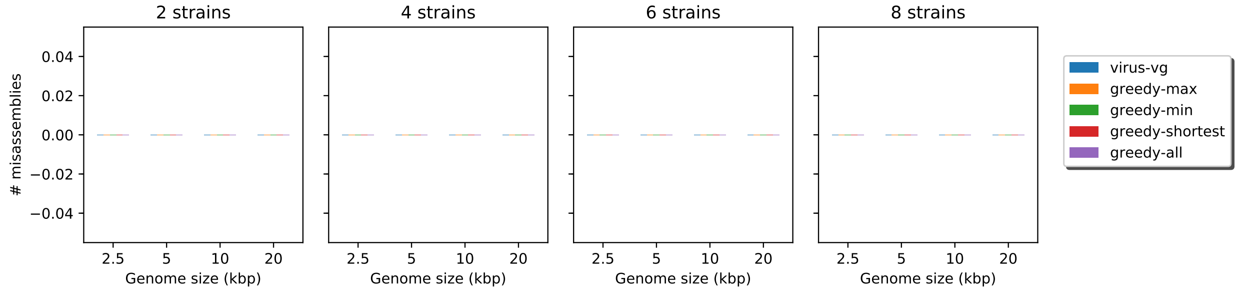

(g) # misassemblies

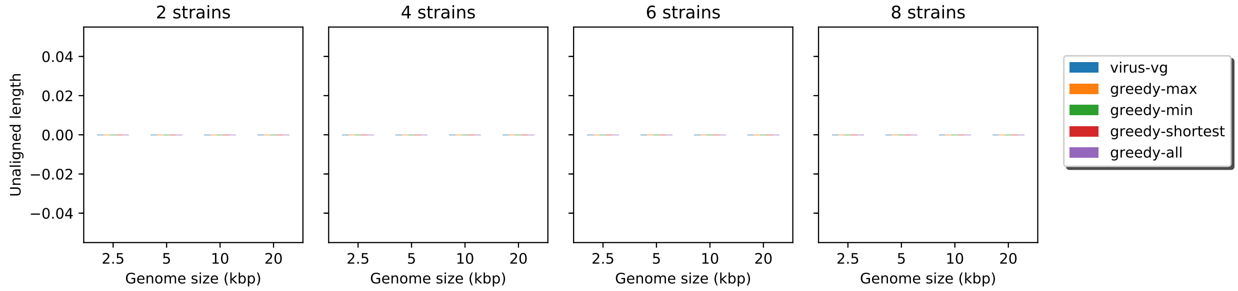

(h) Unaligned length

Figure 10 (cont.): Assembly statistics for data sets of increasing genome size (2500 bp, 5000 bp, 10.000 bp, 20.000 bp) and increasing number of strains (2, 4, 6, 8). Virus-VG results are missing for samples of 6 and 8 strains for genomes of size 5000 bp and up, as well as for samples of 4 strains for genomes of size at least 10.000 bp, because path enumeration became infeasible. Note: there are no misassemblies or unaligned contigs.

### 7 Command lines

For benchmarking purposes we made use of several existing tools, listed below. All tools were run using default settings unless specified otherwise.

*Read simulations:*
SimSeq [26]:
```
java -jar SimSeq-master/SimSeq.jar -l 600 -1 250 -2 250 \
-e SimSeq-master/profiles/miseq_250bp.txt -r truth.fasta \
-n <num_reads> -o sim_reads.sam
```

*Read trimming, adapter removal:*
CutAdapt [27]:
```
cutadapt -b TACACTCTTTCCCTACACGACGCTCTTCCGATCT \
-B GTGACTGGAGTTCAGACGTGTGCTCTTCCGATCT -q 20 -o forward.trimmed.fastq \
-p reverse.trimmed.fastq forward.fastq reverse.fastq
```

*Ad-hoc reference construction:*
VICUNA [28]:
```
vicuna-omp-v1.0 config.txt
```

*Alignment:*
BWA-MEM [29]:
```
bwa mem reference.fasta reads.fastq > reads.sam
```

*Reference-guided quasispecies reconstruction tools:*
ShoRAH [5]:
```
python shorah.py -b paired.sorted.bam -f reference.fasta
```
PredictHaplo [4]:
```
PredictHaplo-Paired config.txt
```

*De novo assembly tools:*
SAVAGE [8]:
```
savage -p1 forward.fastq -p2 reverse.fastq --revcomp --split 30 -t 8
```
Virus-VG [9]:
```
python build_graph_msga.py -f forward.fastq -r reverse.fastq \
    -c contigs_stage_c.fasta -t 8 -vg vg-v1.7.0
python optimize_strains.py -m 100 -c 200 node_abundance.txt \
        contig_graph.final.gfa
```
SPAdes [18]:
```
spades.py -1 forward.fastq -2 reverse.fastq --careful -t 8 -o .
```
VG-flow:
```
python vg-flow.py -m 100 -c 200 --greedy_mode=all \
node_abundance.txt contig_graph.final.gfa
```

*Assembly evaluation:*
QUAST [25]:
```
python quast.py -m 500 -R ground_truth.fasta contigs.fasta
```

### References

[1] M.S. Lindner and B.Y. Renard. Metagenomic abundance estimation and diagnostic testing on species level. *Nucleic Acids Research*, 41(1):e10, 2012.

[2] M. Fischer, B. Strauch, and B.Y. Renard. Abundance estimation and differential testing on strain level in metagenomics data. *Bioinformatics*, 33(14):i124–i132, 2017.

[3] N.L. Bray, H. Pimentel, P. Melsted, and L. Pachter. Near-optimal probabilistic RNA-seq quantification. *Nature Biotechnology*, 34:525–527, 2016.

[4] S. Prabhakaran, M. Rey, O. Zagordi, N. Beerenwinkel, and V. Roth. HIV haplotype inference

using a propagating dirichlet process mixture model. *IEEE Transactions on Computational Biology and Bioinformatics*, 11(1):182–191, 2014.

[5] O. Zagordi, A. Bhattacharya, N. Eriksson, and N. Beerenwinkel. ShoRAH: estimating the genetic diversity of a mixed sample from next-generation sequencing data. *BMC Bioinformatics*, 12(1):119, 2011.

[6] M.C.F. Prosperi and M. Salemi. QuRe: software for viral quasispecies reconstruction from next-generation sequencing data. *Bioinformatics*, 28(1):132–133, Jan 2012.

[7] S. Ahn and H. Vikalo. aBayesQR: A bayesian method for reconstruction of viral populations characterized by low diversity. *Journal of Computational Biology*, 25(7):637–648, 2018.

[8] J.A. Baaijens, A. Zine El Aabidine, E. Rivals, and A Schönhuth. De novo assembly of viral quasispecies using overlap graphs. *Genome Research*, 27(5):835–848, 2017.

[9] J.A. Baaijens, B. Van der Roest, J. Köster, L. Stougie, and A. Schönhuth. Full-length de novo viral quasispecies assembly through variation graph construction. *Bioinformatics*, 05 2019. btz443.

[10] S. Barik, S. Das, and H. Vikalo. Qsdpr: Viral quasispecies reconstruction via correlation clustering. *Genomics*, 110(6):375 – 381, 2018.

[11] J. Chen, Y. Zhao, and Y. Sun. De novo haplotype reconstruction in viral quasispecies using paired-end read guided path finding. *Bioinformatics*, 34(17):2927–2935, 2018.

[12] S. Knyazev, V. Tsyvina, A. Melnyk, A. Artyomenko, T. Malygina, Y.B. Porozov, E. Campbell, W.M. Switzer, P. Skums, and A. Zelikovsky. CliqueSNV: Scalable reconstruction of intra-host viral populations from NGS reads. bioRxiv:10.1101/264242, 2018.

[13] A.I. Tomescu, A. Kuosmanen, R. Rizzi, and V. Mäkinen. A novel min-cost flow method for estimating transcript expression with RNA-seq. *BMC Bioinformatics*, 14(5):S15, Apr 2013.

[14] R. Rizzi, A.I. Tomescu, and V. Mäkinen. On the complexity of minimum path cover with subpath constraints for multi-assembly. *BMC Bioinformatics*, 15(9):S5, 2014.

[15] E. Bernard, L. Jacob, J. Mairal, and J. Vert. Efficient RNA isoform identification and quantification from RNA-seq data with network flows. *Bioinformatics*, 30(17):2447–2455, 2014.

[16] M. Pertea, G.M. Pertea, C.M. Antonescu, T. Chang, J.T. Mendell, and S.L. Salzberg. StringTie enables improved reconstruction of a transcriptome from RNA-seq reads. *Nature Biotechnology*, 33:290–295, 2015.

[17] C. Trapnell, B.A. Williams, G. Pertea, A. Mortazavi, G. Kwan, M.J. van Baren, S.L. Salzberg, B.J. Wold, and L. Pachter. Transcript assembly and quantification by RNA-seq reveals unannotated transcripts and isoform switching during cell differentiation. *Nature Biotechnology*, 28:511–515, 2010.

[18] A. Bankevich, S. Nurk, D. Antipov, A.A. Gurevich, M. Dvorkin, A.S. Kulikov, V.M. Lesin, S.I. Nikolenko, S. Pham, A.D. Prijbelski, A.V. Pyshkin, A.V. Sirotkin, N. Vyahni, G. Tesler, P.A. Pevzner, and M.A. Alekseyev. SPAdes: A new genome assembly algorithm and its applications to single-cell sequencing. *Journal of Computational Biology*, 19(5):455–477, 2012.

[19] A. Sczyrba, Peter Hofmann, Peter Belmann, David Koslicki, Stefan Janssen, Johannes Dröge, Ivan Gregor, Stephan Majda, and A. McHardy. Critical assessment of metagenome interpretation - a benchmark of metagenomics software. *Nature Methods*, 14:1063–1071, 2017.

[20] S. Nurk, D. Meleshko, A. Korobeynikov, and P.A. Pevzner. metaSPAdes: a new versatile metagenomic assembler. *Genome Research*, 27(5):824–834, 2017.

[21] S. Boisvert, F. Raymond, E. Godzaridis, F. Laviolette, and J. Corbeil. Ray meta: scalable de novo metagenome assembly and profiling. *Genome Biology*, 13(12):R122, 2012.

[22] Y. Peng, H.C. Leung, S.M. Yiu, and F.Y. Chin. Meta-IDBA: a de novo assembler for metagenomic data. *Bioinformatics*, 27(13):i94–i101, 2012.

[23] Dinghua Li, Chi-Man Liu, Ruibang Luo, Kunihiko Sadakane, and Tak-Wah Lam. MEGAHIT: an ultra-fast single-node solution for large and complex metagenomics assembly via succinct de Bruijn graph. *Bioinformatics*, 31(10):1674–1676, 01 2015.

[24] F. Di Giallonardo, A. Töpfer, M. Rey, S. Prabhakaran, Y. Duport, C. Leemann, S. Schmutz, N.K. Campbell, B. Joos, M.R. Lecca, A. Patrignani, M. Däumer, C. Beisel, P. Rusert, A. Trkola, H.F. Günthard, V. Roth, N. Beerenwinkel, and K.J. Metzner. Full-length haplotype reconstruction to infer the structure of heterogeneous virus populations. *Nucleic Acids Research*, 42:e115, 2014.

[25] A. Gurevich, V. Saveliev, N. Vyahhi, and G. Tesler. QUAST: quality assessment tool for genome assemblies. *Bioinformatics*, 29(8):1072–1075, 2013.

[26] John St. John. An illumina paired-end and mate-pair short read simulator. https://github.com/jstjohn/SimSeq, 2014.

[27] Marcel Martin. Cutadapt removes adapter sequences from high-throughput sequencing reads. *EMBnet.journal*, 17(1):10–12, 2011.

[28] X. Yang, P. Charlebois, S. Gnerre, M. Coole, N. Lennon, J. Levin, J. Qu, E. Ryan, M. Zody, and M. Henn. De novo assembly of highly diverse viral populations. *BMC Genomics*, 13(1):475, 2012.

[29] H. Li. Aligning sequence reads, clone sequences and assembly contigs with BWA-MEM. arXiv:1303.3997, 2013.
